# Gene expression in African Americans and Latinos reveals ancestry-specific patterns of genetic architecture

**DOI:** 10.1101/2021.08.19.456901

**Authors:** Linda Kachuri, Angel C.Y. Mak, Donglei Hu, Celeste Eng, Scott Huntsman, Jennifer R. Elhawary, Namrata Gupta, Stacey Gabriel, Shujie Xiao, Kevin L. Keys, Akinyemi Oni-Orisan, José R. Rodríguez-Santana, Michael LeNoir, Luisa N. Borrell, Noah A. Zaitlen, L. Keoki Williams, Christopher R. Gignoux, Esteban González Burchard, Elad Ziv

**Affiliations:** Department of Epidemiology & Biostatistics, University of California San Francisco, San Francisco, CA, USA; Department of Medicine, University of California San Francisco, San Francisco, CA, USA; Broad Institute of MIT and Harvard, Cambridge, MA, USA; Center for Individualized and Genomic Medicine Research, Henry Ford Health System, Detroit, MI, USA; Berkeley Institute for Data Science, University of California, Berkeley, CA, USA; Department of Clinical Pharmacy, University of California, San Francisco, San Francisco, CA, USA; Department of Bioengineering and Therapeutic Sciences, University of California San Francisco, San Francisco, CA, USA; Institute for Human Genetics, University of California San Francisco, San Francisco, CA, USA; Centro de Neumología Pediátrica, San Juan, Puerto Rico; Bay Area Pediatrics, Oakland, CA, USA; Department of Epidemiology and Biostatistics, Graduate School of Public Health and Health Policy, City University of New York, New York, NY, USA; Department of Neurology, University of California, Los Angeles, Los Angeles, CA, USA; Department of Computational Medicine, University of California, Los Angeles, Los Angeles, CA, USA; Department of Internal Medicine, Henry Ford Health System, Detroit, MI, USA; Colorado Center for Personalized Medicine, University of Colorado Anschutz Medical Campus, Aurora, CO, USA; Department of Biostatistics and Informatics, School of Public Health, University of Colorado Anschutz Medical Campus, Aurora, CO, USA; Helen Diller Family Comprehensive Cancer Center, University of California San Francisco, San Francisco, CA, USA

## Abstract

We analyzed whole genome and RNA sequencing data from 2,733 African American and Hispanic/Latino children to explore ancestry- and heterozygosity-related differences in the genetic architecture of whole blood gene expression. We found that heritability of gene expression significantly increases with greater proportion of African genetic ancestry and decreases with higher levels of Indigenous American ancestry, consistent with a relationship between heterozygosity and genetic variance. Among heritable protein-coding genes, the prevalence of statistically significant ancestry-specific expression quantitative trait loci (anc-eQTLs) was 30% in African ancestry and 8% for Indigenous American ancestry segments. Most of the anc-eQTLs (89%) were driven by population differences in allele frequency, demonstrating the importance of measuring gene expression across multiple populations. Transcriptome-wide association analyses of multi-ancestry summary statistics for 28 traits identified 79% more gene-trait pairs using models trained in our admixed population than models trained in GTEx. Our study highlights the importance of large and ancestrally diverse genomic studies for enabling new discoveries of complex trait architecture and reducing disparities.

## INTRODUCTION

Gene expression has been extensively studied as a trait affected by genetic variation in humans^1^. Expression quantitative trait loci (eQTLs) have been identified in most genes^2–4^ and extensive analyses across multiple tissues have demonstrated both tissue-specific and shared eQTLs^2^. Genome-wide association studies (GWAS) tend to identify loci that are enriched for eQTLs^5^. Colocalization of eQTLs with GWAS has become an important element of identifying causal genes and investigating the biology underlying genetic susceptibility to disease^6^. More recently, transcriptome-wide association studies (TWAS) have been developed to systematically leverage eQTL data by imputing transcriptomic profiles in external datasets, which has led to the discovery of trait-associated genes that were often missed by GWAS^7, 8^.

GWAS have identified thousands of loci for hundreds of diseases and disease-related phenotypes in human populations^9^. However, non-European ancestry populations are significantly under-represented in GWAS^10, 11^ and in studies of gene expression and eQTLs. We and others have shown that gene expression prediction models trained in predominantly European ancestry reference datasets, such as the Genotype-Tissue Expression (GTEx) project^2^, have substantially lower accuracy to predict gene expression levels when applied to populations of non-European ancestry^3, 12, 13^. The importance of having ancestry-matched training datasets for prediction accuracy is also reflected by the limited cross-population portability of other multi-SNP prediction models, such as polygenic risk scores (PRS)^14–16^. Therefore, the limited diversity in genetic association studies and reference datasets is a major obstacle for applying existing integrative genomic studies to non-European populations.

To address this gap, we leveraged whole genome and RNA sequencing data from 2,733 African American and Latino children from the Genes-environments and Admixture in Latino Americans (GALA II) study and the Study of African Americans, Asthma, Genes, and Environments (SAGE) to characterize the genetic architecture of whole blood eQTLs. The diversity within the GALA II/SAGE population enabled us to evaluate how genetic ancestry relates to the heritability of gene expression, and systematically quantify the prevalence of ancestry-specific eQTLs. Lastly, we developed a powerful set of TWAS models from these datasets to facilitate genetic association analyses in multi-ancestry populations.

## RESULTS

### Demographic characteristics of GALA II and SAGE participants

We analyzed data from a total of 2,733 participants from the GALA II and SAGE asthma case-control studies, including 757 self-identified African Americans (AA), 893 Puerto Ricans (PR), 784 Mexican Americans (MX), and 299 other Latinos (LA) who did not self-identify as Mexican American or Puerto Rican (Table 1, Table S1). All four grandparents of each study participant also identified as Latino for GALA II or African American for SAGE. The median age of the participants varied from 13.2 (PR) to 16.0 (AA) years old. About 50% of the participants were female and 45% (MX) to 62% (PR) had physician-diagnosed asthma. For each participant we estimated genome-wide genetic ancestry (global ancestry) proportions, visualized in Figure 1. Median global African ancestry was highest in AA (82.6%), followed by PR (19.7%), and lowest in MX (3.5%).

**Figure 1.**
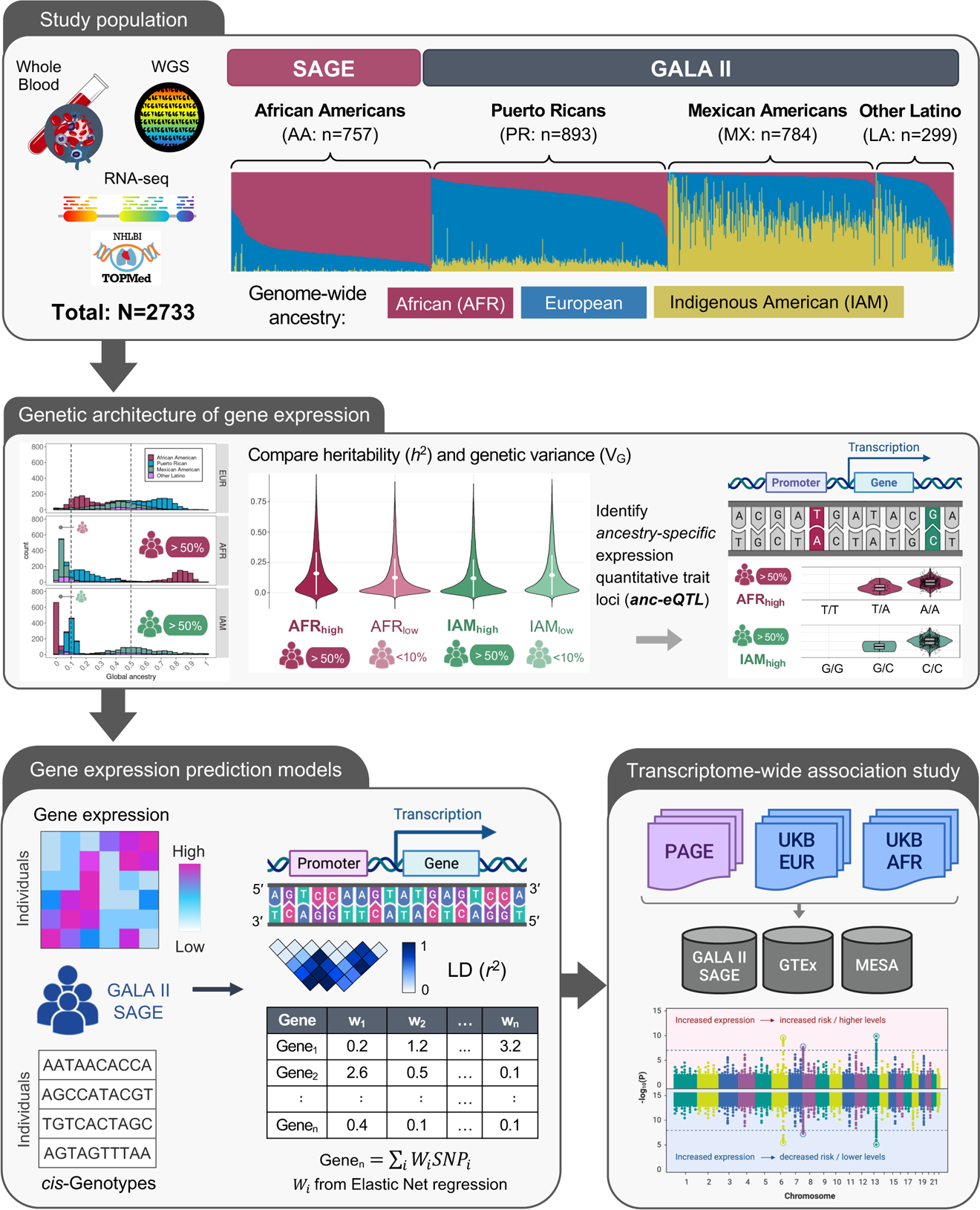
Study Overview. This study included TOPMed whole genome sequencing and whole transcriptome data generated from whole blood samples of SAGE African American and GALA II Latino individuals (n=2,733). We compared elements of the genetic architecture gene expression, such as and *cis*-heritability and genetic variance, across participant groups defined by self-identified race/ethnicity and genetic ancestry. Next, we developed genetic prediction models of whole blood transcriptome levels and performed comparative transcriptome-wide association studies (TWAS) using GWAS summary statistics generated from the PAGE study and the UK Biobank.

**Table 1:** Study Participants. Demographic characteristics of 2,733 participants from the Genes-environments and Admixture in Latino Americans (GALA II) and the Study of African Americans, Asthma, Genes, and Environments (SAGE) included in the present analysis.

|  | Self-identified Race/Ethnicity |  |  |  |  |  |  |  | Pooled |  |
| --- | --- | --- | --- | --- | --- | --- | --- | --- | --- | --- |
|  | African American |  | Puerto Rican |  | Mexican |  | Other Latino |  |  |  |
|  | N | (%) | N | (%) | N | (%) | N | (%) | N | (%) |
| **Sex** |  |  |  |  |  |  |  |  |  |  |
| Female | 405 | (53.5) | 451 | (50.5) | 427 | (54.5) | 158 | (52.8) | 1,441 | (52.7) |
| **Asthma status** |  |  |  |  |  |  |  |  |  |  |
| Case | 433 | (57.2) | 549 | (61.5) | 351 | (44.8) | 156 | (52.2) | 1489 | (54.5) |
| **Recruitment center** |  |  |  |  |  |  |  |  |  |  |
| SF Bay Area | 757 | (100) | 0 | (0) | 348 | (44.4) | 109 | (36.5) | 1214 | (44.4) |
| Chicago | 0 | (0) | 31 | (3.5) | 247 | (31.5) | 52 | (17.4) | 330 | (12.1) |
| Puerto Rico | 0 | (0) | 837 | (93.7) | 0 | (0) | 8 | (2.7) | 845 | (30.9) |
| New York City | 0 | (0) | 22 | (2.5) | 36 | (4.6) | 86 | (28.8) | 144 | (5.3) |
| Houston | 0 | (0) | 3 | (0.3) | 153 | (19.5) | 44 | (14.7) | 200 | (7.3) |
|  | **Median** | **(IQR)** | **Median** | **(IQR)** | **Median** | **(IQR)** | **Median** | **(IQR)** | **Median** | **(IQR)** |
| **Age (years)** | 16.0 | (6.6) | 13.2 | (4.8) | 13.8 | (6.5) | 13.7 | (5.7) | 14.0 | (6.3) |
| **Genetic ancestry (%)** |  |  |  |  |  |  |  |  |  |  |
| African | 82.6 | (9.4) | 19.7 | (13.3) | 3.5 | (2.7) | 8.3 | (14.8) | 17.5 | (61.8) |
| Indigenous American | 0.3 | (0.9) | 9.9 | (3.6) | 55.3 | (23.2) | 42.3 | (43.2) | 10.7 | (45.2) |
| European | 16.5 | (9.5) | 69.5 | (13.6) | 40.3 | (21.9) | 45.9 | (20.8) | 44.2 | (43.8) |
| **Total** | **757** |  | **893** |  | **784** |  | **299** |  | **2733** |  |

Abbreviations
IQR Interquartile range

### Variability in gene expression accounted by common genetic variation increases with African ancestry

We compared the heritability (h^2^) and genetic variance (V_G_) of whole blood gene expression across self-identified race/ethnicity groups (AA, PR, MX) and populations defined based on genetic ancestry. There was a positive association between increasing proportion of African ancestry and variability of gene expression attributed to common genetic variation (minor allele frequency [MAF] ≥0.01) within the *cis*-region (see Methods). Across 17,657 genes, *cis*-heritability (Figure 2A) was significantly higher in AA (median h^2^=0.097) compared to PR (h^2^=0.072; Wilcoxon rank sum test: p=2.2×10^-50^) and MX (h^2^=0.059; p=3.3×10^-134^), as well as PR compared to MX (p=2.2×10^-25^). Genetic variance (Figure 2B) of whole blood transcript levels in AA (median V_G_=0.022) was higher than in PR (V_G_=0.018, p=4.0×10^-19^) and in MX (V_G_=0.013, p=5.6×10^-135^). Results remained unchanged when sample size was fixed to n=600 in all populations (Figure S1), with higher heritability and genetic variance in AA (h^2^=0.098; V_G_=0.022) compared to PR (h^2^=0.072; V_G_=0.017) and MX (h^2^=0.062; V_G_=0.012).

**Figure 2.**
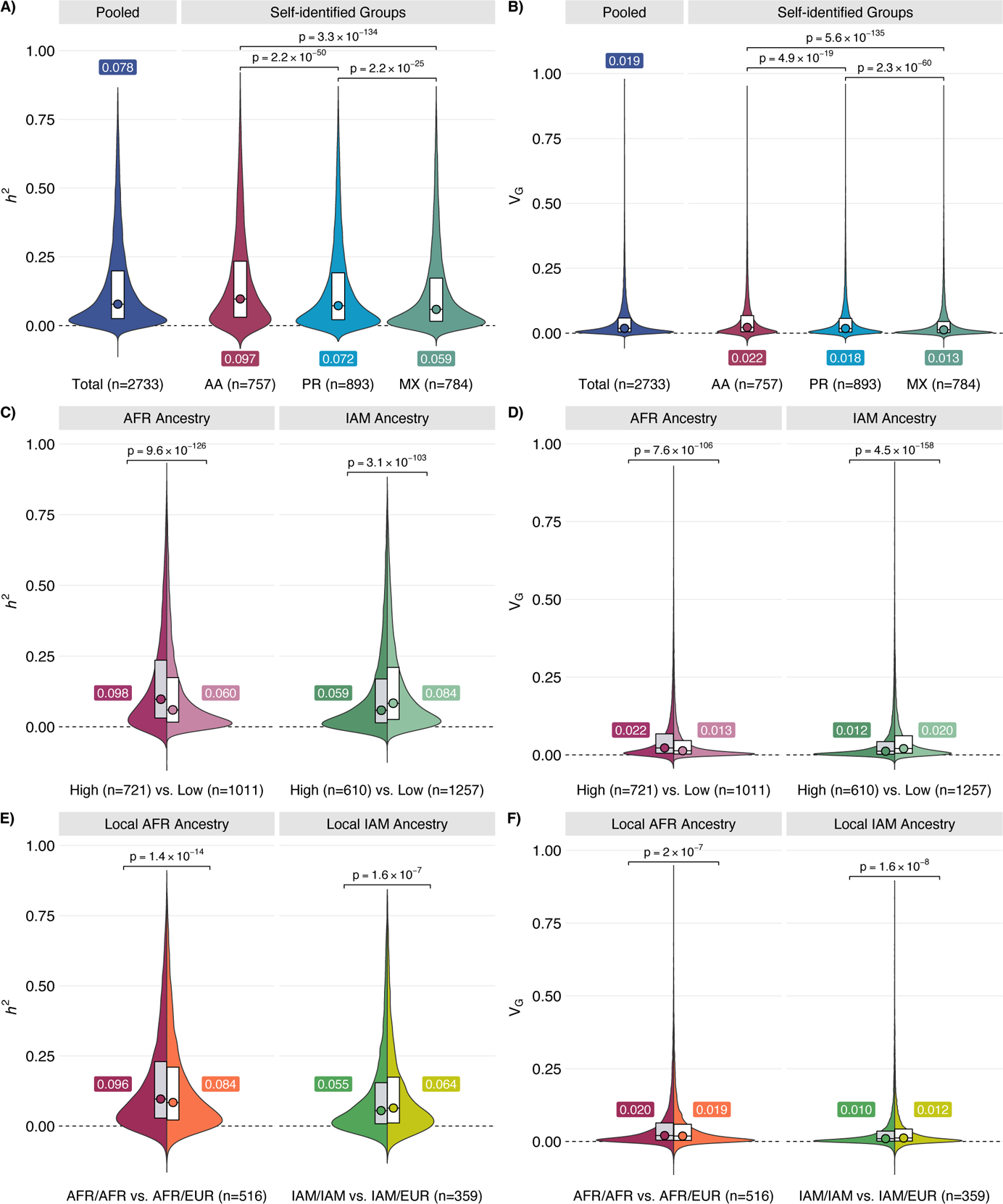
Comparison of *cis*-heritability (*h^2^*) and genetic component of transcriptome variance (V_G_) by self-identified race/ethnicity and genetic ancestry groups. Analyses stratified by self-identified race/ethnicity **(A-B)** and genetic ancestry comparing individuals with >50% global ancestry (High) to participants with <10% of the same ancestry (Low) **(C-D)**. Local ancestry at the transcriptional start site of each gene was used to compare subjects with 100% (AFR/AFR or IAM/IAM) to 50% (AFR/EUR or IAM/EUR) local ancestry **(E-F)**. Median values of h^2^ or V_G_ and two-sided Wilcoxon p-values are annotated.

Next, we compared the distribution of h^2^ (Figure 2C) and V_G_ (Figure 2D) between participants grouped based on proportions of global genetic ancestry (Table S3). Among participants with >50% African ancestry (AFR_high_, n=721) *cis-*heritability (h^2^=0.098) and genetic variance (V_G_=0.022) were higher than in n=1011 participants with <10% global African ancestry (AFR_low_: h^2^=0.060, *P*_Wilcoxon_=9.6×10^-126^; V_G_=0.013, *P*_Wilcoxon_=7.6×10^-106^). Among individuals with >50% Indigenous American (IAM) ancestry (IAM_high_, n=610), *cis*-heritability (h^2^=0.059) and genetic variance (V_G_=0.012) were lower than in subjects with <10% IAM ancestry (IAM_low_: h2=0.084, p=3.1×10^-103^; V_G_=0.020, *P*_Wilcoxon_=3.1×10^-158^). To further characterize these findings, we partitioned h^2^ and V_G_ by coarse MAF bins (Figure S2). Although h^2^ and V_G_ remained higher in AFR_high_ compared to AFR_low,_ the magnitude of this difference was more pronounced in the 0.01≤ MAF ≤ 0.10 bin (h^2^: 0.032 vs. 0.013, *P*_Wilcoxon_=1.8×10^-310^) than for variants with MAF>0.10 (h^2^: 0.038 vs. 0.027, *P*_Wilcoxon_=2.2×10^-55^). Larger differences in h^2^ and V_G_ among 0.01≤ MAF ≤ 0.10 variants were also observed for IAM_high_ and IAM_low_.

We also investigated the impact of ancestry at the locus level, defined as the number of alleles (0, 1 or 2) derived from each ancestral population at the transcription start site (Table S4). For each gene, individuals with homozygous local African ancestry (AFR/AFR) were compared to those with heterozygous local African and European ancestry (AFR/EUR). Heritability was significantly higher in AFR/AFR homozygotes (h^2^=0.096) compared to AFR/EUR (h^2^=0.084, *P*_Wilcoxon_=1.4×10^-14^), and lower in IAM/IAM (h^2^=0.055) compared to IAM/EUR (h^2^=0.064, p=1.6×10^-7^; Figure 2E). Compared to global ancestry, the magnitude of differences in V_G_ was attenuated, but remained statistically significant for AFR (V_G_=0.020 vs. V_G_=0.019, p=2.0×10^-7^) and IAM (V_G_=0.010 vs. V_G_=0.012, p=1.62×10^-8^; Figure 2F; Table S4). Results were also consistent for V_G_ comparisons within race/ethnicity groups for AFR (AA: *P*_Wilcoxon_=5.7×10^-5^; PR: *P*_Wilcoxon_=2.0×10^-7^) and IAM (MX: p=2.0×10^-7^) (Table S4).

As a parallel approach to evaluating heritability, we applied LDAK (Linkage Disequilibrium Adjusted Kinships), which assumes that SNP-specific variance is inversely proportional not only to MAF, but also to LD tagging^17^. Estimates obtained using LDAK-Thin and GCTA were nearly identical for self-identified groups (AA: h^2^=0.094; PR: h^2^=0.071; MX: h^2^=0.059) and across strata based on global genetic ancestry (AFR_high_: h^2^=0.104; AFRlow: h^2^=0.066, IAM_high_: h^2^=0.062; IAM_low_ h^2^=0.093), suggesting that our results were not sensitive to the assumptions of the GCTA model (Table S5).

Lastly, we tabulated the number of heritable genes for which global and/or local ancestry was significantly associated (FDR<0.05) with transcript levels (Figure S3). Global AFR ancestry was associated with the expression of 326 (2.4%) and 589 (4.5%) of heritable genes in AA and PR, respectively (Table S6). Associations with local, but not global, AFR ancestry were more common (8.9% in AA; 10.9% in PR), and relatively few genes were associated with both measures of ancestry (1.5% in AA and 2.5% in PR). Among genes associated with both global and local AFR ancestry in AA, global AFR ancestry explained 1.8% of variation in gene expression, while local AA accounted for 3.8% (Figure S3). Local IAM ancestry was associated with the expression of 9.8% of genes in MX, compared to 2.8% for global IAM ancestry. Among genes associated with both, local IAM ancestry accounted for 3.5% variation in transcript abundance, while global IAM ancestry accounted for 1.8%.

### Assessment of ancestry-specific eQTLs

We next sought to understand patterns of cis-eQTLs in the admixed GALA/SAGE study participants. A total of 19,567 genes with at least one cis-eQTL (eGenes) were found in the pooled sample. The largest number of eGenes was detected in AA (n=17,336), followed by PR (n=16,975), and MX (n=15,938) participants (Table S7, Figure S4). In analyses stratified by global genetic ancestry the number of eGenes was similar in AFR_high_ (n=17,123) and AFR_low_ (n=17,146) groups (Table S7). When sample size was fixed to n=600 for all ancestry groups (Table S7), the highest number of eGenes (n=16,100) was observed in AFR_high_, followed by IAM_low_ (n=14,866), IAM_high_ (n=14,419), and AFR_low_ (n=14,344). The number of LD-independent (r^2^<0.10) cis-eQTLs per gene was significantly higher in AFR_high_ than AFR_low_ (*P*_Wilcoxon_=2.7×10^-246^), with 63% of genes having more independent cis-eQTLs in AFR_high_ compared to AFR_low_ (Figure S5). Conversely, the number of independent cis-eQTLs detected in IAM_high_ was lower than in IAM_low_ (*P*_Wilcoxon_=2.8×10^-33^).

To characterize ancestry-related differences in the genetic regulation of gene expression, we developed a three-tier framework for identifying ancestry-specific eQTLs, which we refer to as anc-eQTLs (see Methods; Figure 3A; Table S8-S9). For heritable protein-coding genes, we first compared the overlap in 95% credible sets of *cis*-eQTLs identified in participants with >50% global ancestry (AFR_high_; IAM_high_) and those with <10% of the same global ancestry (AFR_low_; IAM_low_). For genes with non-overlapping 95% credible sets, we distinguished between population differences in MAF (Tier 1) and LD (Tier 2). For genes with overlapping 95% credible sets, eQTLs were further examined for evidence of effect size heterogeneity between ancestry groups (Tier 3).

**Figure 3.**
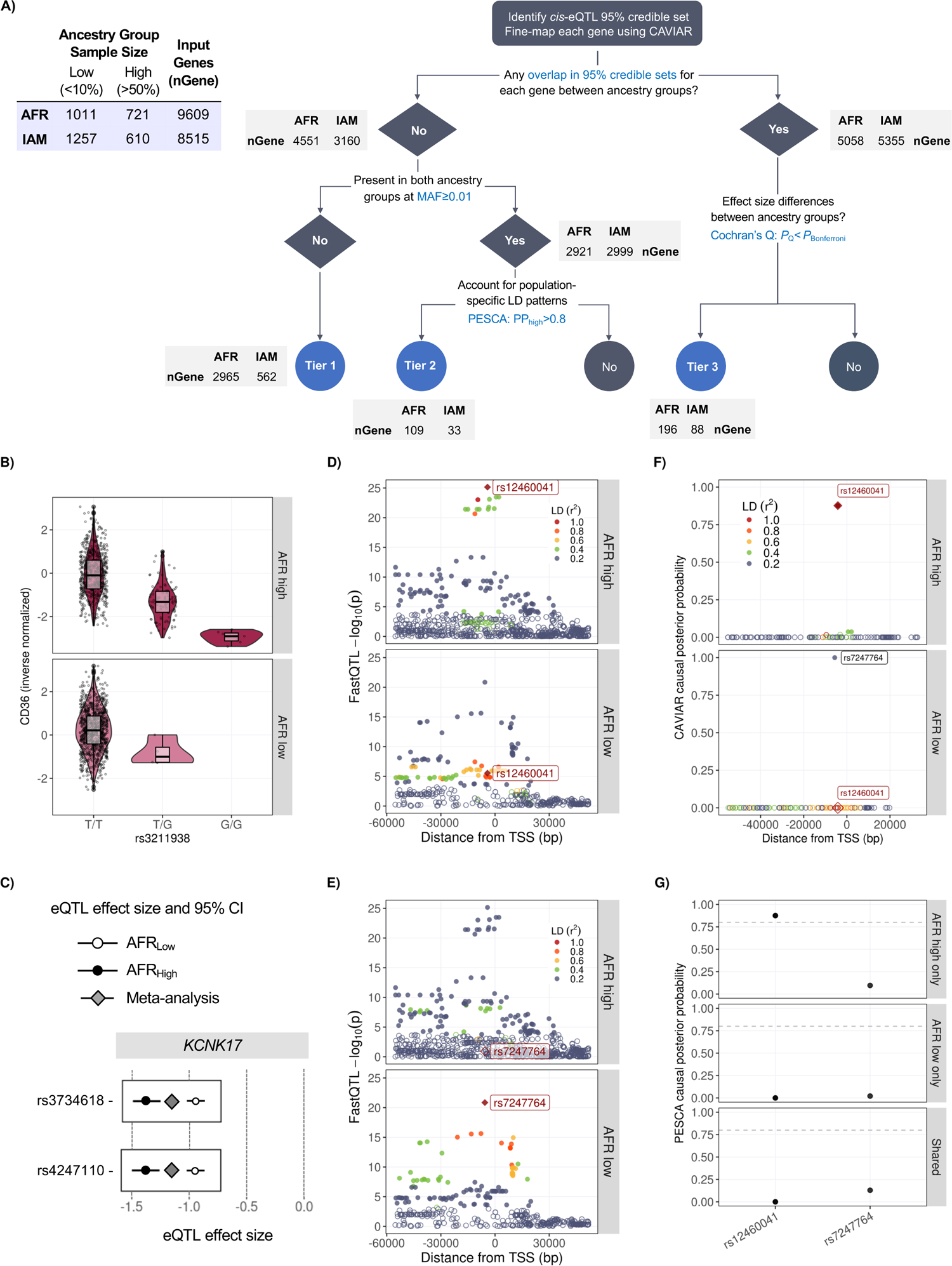
Identification of ancestry-specific eQTLs (anc-eQTLs). **A)** Decision tree for the identification of anc-eQTLs. Number of genes remaining after each step is indicated alongside each branch. **B)** An example of a Tier 1 AFR_high_ anc-eQTL (rs3211938) for *CD36*. **C)** An example of Tier 3 AFR_high_ anc-eQTLs (rs34247110 and rs3734618) for *KCNK17*. Both eQTLs from the 95% credible set had significantly different effect sizes in AFR_high_ and AFR_low_ populations. **D-G)** An example of a Tier 2 AFR_high_ anc-eQTL (rs12460041) for *TRAPPC6A*. CAVIAR detected different lead eQTLs with non-overlapping credible sets in AFR_high_ **(D)** and AFR_low_ **(E)** groups. In each panel variants are colored based on LD r^2^ with respect to index variant (diamond) and eQTLs are denoted by filled circles. **F)** The lead eQTL in AFR_high_ (rs12460041) had a posterior probability (PP)=0 in AFR_low_. **G)** Fine-mapping using PESCA confirmed rs12460041 as a Tier 2 anc-eQTL with PP>0.80 in AFR_high_.

Tier 1 anc-eQTLs (ancestry-specific enrichment) were common (MAF ≥ 0.01) only in individuals with >50% AFR or IAM ancestry and were thus considered to be the most ancestry specific. Over 28% (n=2,695) of genes contained at least one Tier 1 AFR_high_ anc-eQTL, while 7% (n=562) of genes contained a Tier 1 IAM_high_ anc-eQTL (Table S9). A representative example of a Tier 1 AFR_high_ anc-eQTL is rs3211938 (*CD36*), which has MAF=0.077 in AFR_high_ and MAF=0.0020 in AFR_low_, (Figure 3B). This variant been linked to high density lipoprotein (HDL) cholesterol levels in several multi-ancestry GWAS that included African Americans^18–20^.

Tier 2 anc-eQTLs (ancestry-specific LD patterning) had MAF ≥ 0.01 in both high (>50%) and low (<10%) global ancestry groups and were further interrogated using PESCA^21^ to account for population-specific LD patterns. There were 109 genes (1.1%) that contained eQTLs with a posterior probability (PP) >0.80 of being specific to AFR_high_ and 33 genes (0.4%) matching the same criteria for IAM_high_ (Table S9). For instance, two lead eQTLs with non-overlapping credible sets were detected for *TRAPPC6A* in AFR_high_ (rs12460041) and AFR_low_ (rs7247764) groups (Figure 3D-3F). These variants were in low LD (r^2^=0.10 in AFR_high_ and r^2^=0.13 in AFR_low_) and PESCA analysis confirmed that rs12460041 was specific to AFR_high_ (PP>0.80).

Over 50% of heritable protein-coding genes (AFR: n=5,058; IAM: n=5,355) had overlapping 95% credible sets of eQTLs between high and low ancestry groups. Among these shared signals, there was a small proportion of eQTLs that exhibited significant effect size heterogeneity (Tier 3, ancestry-related heterogeneity: 2.0% for AFR_high_; 1.0% for IAM_high_). For instance, rs34247110 and rs3734618 were included in 95% credible sets for *KCNK17* in AFR_high_ and AFR_low_ with significantly different effect sizes (Cochran’s Q p-value=1.8×10^-10^) in each population (Figure 3C). One of these variants, rs34246110, was associated with type 2 diabetes in two independent studies performed in Japanese and multi-ancestry (European, African American, Hispanic and Asian) populations^22, 23^. The detection of this variant in multiple populations is consistent with Tier 3 variants denoting eQTL signals that are shared between ancestries but may have different magnitudes of effect.

The prevalence of any Tier 1, 2, or 3 anc-eQTL was 30% (n=2,961) for AFR ancestry and 8% (n=679) for IAM ancestry. Overall, 3,333 genes had anc-eQTLs for either ancestry. The remaining genes (AFR: n=6,648; IAM: n=7,836) did not contain eQTLs with ancestry-related differences in MAF, LD, or effect size as outlined above. Increasing the global ancestry cut-off to >70% did not have an appreciable impact on anc-eQTLs in AFR_high_ (28.1% overall; 27.3% for Tier 1), but substantially decreased the number of anc-eQTLs in IAM_high_ (3.3% overall; 3.3% Tier 1), likely due to a greater reduction in sample size in this group (n=212 vs. n=610; Table S10). Considering all protein-coding genes (n=13,535) without filtering based on heritability, the prevalence of anc-eQTLs is 22% for AFR_high_, 5% for IAM_high_, and 25% overall. The observation that anc-eQTLs were more common in participants with >50% global AFR ancestry aligns with the higher h^2^ and V_G_ in this population, as well as a greater number of LD-independent cis-eQTLs in AFR_high_ compared to AFR_low_ (Figure S5). Among genes with Tier 1 and Tier 2 anc-eQTLs, 83% had higher h^2^ estimates in AFR_high_ than in AFR_low_, while this was observed in 57% of genes without any ancestry-specific eQTLs 57% (Figure S6).

Despite the limited representation of subjects from diverse ancestries in studies from the NHGRI-EBI GWAS catalog^24^, we detected 70 unique anc-eQTLs associated with 84 phenotypes (Table S11). Most of these were Tier 3 anc-eQTLs (59%) that mapped to blood cell traits, lipids, and blood protein levels. To further explore the relevance of the eQTLs identified in our analysis to other complex traits, we performed colocalization with summary statistics for 28 traits from the multi-ancestry PAGE study^20^ (see Methods). We identified 78 eQTL-trait pairs (85 eGene-trait pairs) with strong evidence of a shared genetic, defined as PP_4_>0.80, 16 of which were anc-eQTLs (Table S12). One compelling example is rs7200153, AFR_high_ Tier 1 anc-eQTL for the haptoglobin (*HP*) gene, which colocalized with total cholesterol (PP_4_=0.997; Figure S7). Fine-mapping limited the 95% credible set to two variants in high LD (r^2^=0.75): rs7200153 (PP_SNP_=0.519) and rs5471 (PP_SNP_=0.481). Although rs7200153 had a slightly higher PP_SNP_, rs5471 is likely to be the true causal variant given its proximity to the *HP* promoter, stronger effect of *HP* expression, and experimental data demonstrating decreased transcriptional activity for rs5471-C in West African populations^25–27^. Prior studies have identified *HP* as having an effect on cholesterol and the association of rs5471 is well supported by multi-ancestry genetic association studies^19, 28, 29^.

Although our primary assessment of ancestry-specific eQTLs focused on variants in *cis*, we also performed trans-eQTL analyses that identified 33 trans-eGenes in AA, 52 trans-eGenes in PR, and 51 trans-eGenes in MX subjects (see Methods; Table S13). Analyses stratified by genetic ancestry detected 36 independent (LD r^2^<0.10) trans-eQTLs and 31 eGenes, 26 of which (24 eGenes) were found in AFR_high_ but not in AFR_low_. Fewer independent signals were detected in participants with >50% Indigenous American ancestry (26 trans-eQTLs), of which 23 trans-eQTLs were not detected in the IAM_low_ group.

### Gene expression prediction models from admixed populations increase power for gene discovery

We generated gene expression imputation models from GALA II and SAGE following the PrediXcan approach^7^. We used the pooled population (n=2,733) to generate models with significant prediction (see Methods) for 11,830 heritable genes with mean cross-validation (CV) *R*^2^=0.157 (Table S13, Figure S8). We also generated population-specific models for African Americans (10,090 genes, CV *R*^2^=0.180), Puerto Ricans (9,611 genes, CV *R*^2^=0.163), and Mexican Americans (9,084 genes, CV *R*^2^=0.167). In sensitivity analyses that adjusted for local ancestry (Table S14), we did not observe gains in predictive performance (AA: CV *R*^2^=0.177; PR: CV *R*^2^=0.154; MX: CV *R*^2^=0.159).

Validation of GALA/SAGE TWAS models and comparison with GTEx v8 was performed in the Study of Asthma Phenotypes and Pharmacogenomic Interactions by Race-Ethnicity (SAPPHIRE)^30^, an independent adult population of 598 African Americans (Figure S9). Validation accuracy was proportional to the degree of alignment in ancestry between training and testing study samples. For 5,254 genes with TWAS models available in GALA/SAGE and GTEx, median correlation between genetically predicted and observed transcript levels in SAPPHIRE was highest for pooled (Pearson’s r = 0.086) and AA (Pearson’s r = 0.083) models and lowest for GTEx (Pearson’s r = 0.049).

To evaluate the potential of TWAS models generated in the pooled GALA II and SAGE population (hereafter referred to as GALA/SAGE models) to improve gene discovery in admixed populations, we applied our models to GWAS summary statistics for 28 traits from the multi-ancestry Population Architecture using Genomics and Epidemiology (PAGE) study^20^ and conducted parallel analyses using TWAS models based on GTEx v8^2,7^ and the Multi-Ethnic Study of Atherosclerosis (MESA)^3^. GTEx v8 whole blood models are based on 670 subjects of predominantly European ancestry (85%)^2^. MESA models impute monocyte gene expression^3^ based on a sample of African American and Hispanic/Latino individuals (MESA_AFHI_: n=585). As such, populations included in MESA and PAGE more closely resemble the ancestry composition of our GALA/SAGE populations.

The number of genes with available TWAS models was 39% to 82% higher in GALA/SAGE compared to GTEx (n=7,249) and MESA_AFHI_ (n=5,555). Restricting to 3,143 genes shared across all three models, CV R^2^ was significantly higher in GALA/SAGE compared to GTEx (*P*_Wilcoxon_=4.6×10^-1^^59^) and MESA_AFHI_ (*P*_Wilcoxon_=1.1×10^-64^), which is expected based on the large sample size of GALA/SAGE (Figure 4A). TWAS models generated in GALA/SAGE AA (n=757) attained higher CV R^2^ than GTEx (*P*_Wilcoxon_=2.2×10^-103^), which had a comparable training sample size (n=670), and MESA_AFA_ models (p=6.2×10^-43^) trained in 233 individuals (Figure 4B).

**Figure 4.**
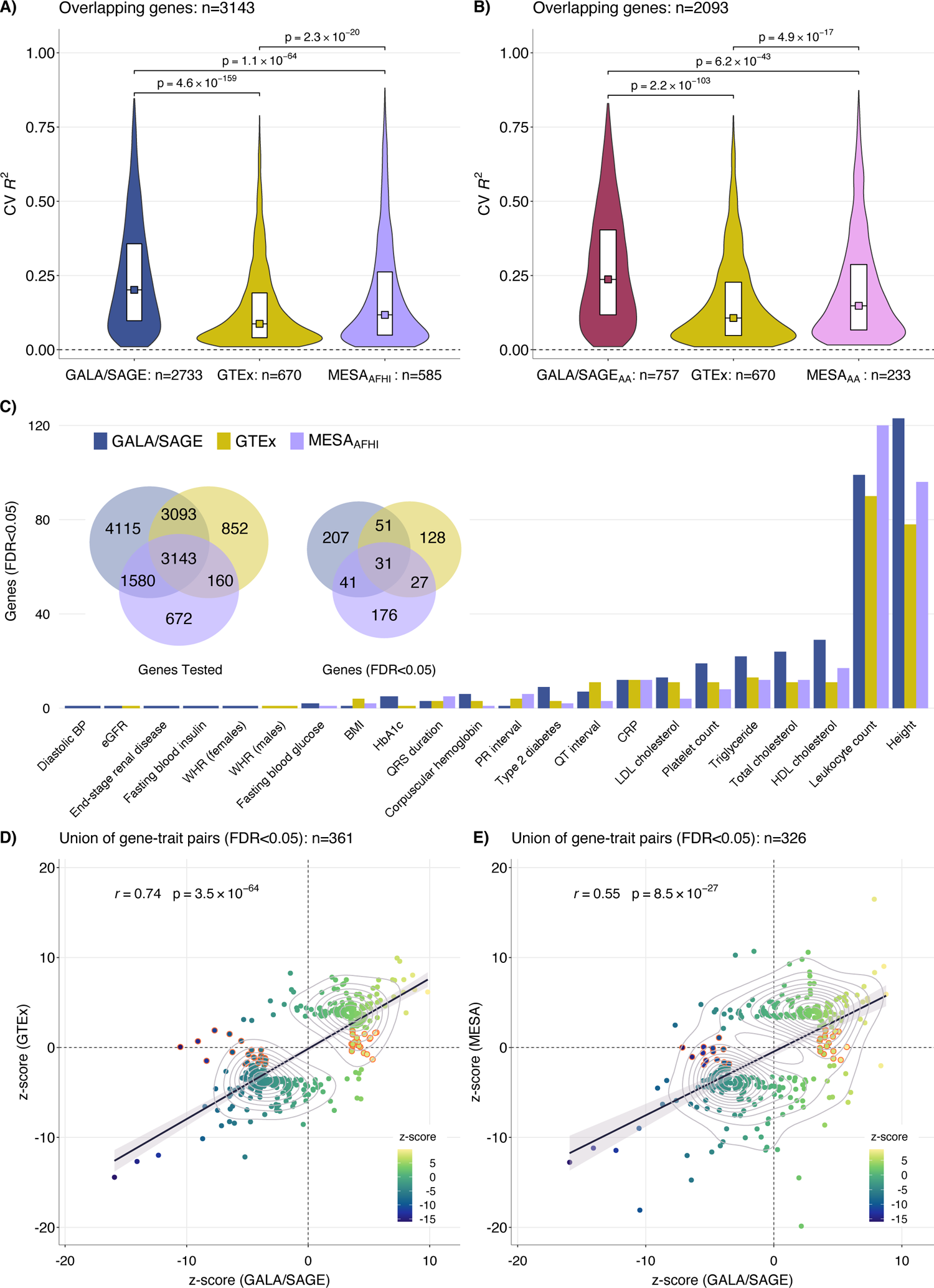
Transcriptome imputation model performance and TWAS results in PAGE. Internal cross-validation R^2^ values for each model were compared for overlapping genes using a two-sided Wilcoxon test. GTEx v8 whole blood TWAS models were compared to models trained in **A)** pooled African American and Hispanic/Latino samples and **B)** African Americans only from GALA/SAGE and MESA, respectively. **C)** Summary of TWAS results for 28 traits in PAGE. Correlation between TWAS z-scores from analyses using GALA/SAGE pooled models and z-scores using **D)** GTEx and **E)** MESA for the union of genes that achieved FDR<0.05 using either prediction model. Genes highlighted in orange had FDR<0.05 using GALA/SAGE models but did not reach nominal significance (TWAS p-value>0.05) using GTEx or MESA models.

Association results across 28 PAGE traits demonstrate that TWAS using GALA/SAGE pooled models identified a larger number of significant gene-trait pairs (n=380, FDR<0.05), followed by MESA_AFHI_ (n=303), and GTEx (n=268), with only 30 genes (35 gene-trait pairs) significant in all three analyses (Figure 4C). GALA/SAGE models yielded a larger number of associated genes than MESA in 80% of analyses (binomial test: p=0.012) and 79% compared to GTEx (binomial test: p=0.019). Of the 330 genes with FDR<0.05 in GALA/SAGE, 143 (43%) were not present in GTEx and 199 (60%) were not present in MESA_AFHI_. For genes that were significant in at least one TWAS, z-scores in GALA/SAGE were highly correlated with GTEx (Figure 4C; *r* =0.74, p=3.5×10^-64^) and MESA_AFHI_ (Figure 4D; *r* = 0.55, p=8.5×10^-27^), suggesting that most genes have concordant effects even if they fail to achieve statistical significance in both analyses. Despite the higher correlation with GTEx z-scores, we observed a higher proportion of gene-trait pairs with FDR<0.05 in GALA/SAGE but not even nominally associated (*P*_TWAS_<0.05) in GTEx (33%), compared to 18% in MESA_AFHI_.

HDL cholesterol exhibited one of the largest differences in TWAS associations, with over 60% more significant genes identified using GALA/SAGE models (n=29) than GTEx predictions (n=11; Figure 4C). TWAS models for several associated genes, including those with established effects on cholesterol transport and metabolism, like *CETP*, were not available in GTEx. The top HDL-associated gene, *CD36* (z-score= −10.52, *P*_TWAS_=6.9×10^-26^) had Tier 1 AFR_high_ anc-eQTLs (rs3211938) that were not present at an appreciable frequency in populations with low African ancestry (MAF in European = 1.3×10^-4^). The difference in MAF may explain why *CD36* was not detected using GTEx (z-score=0.057, *P*_TWAS_=0.95), even though all 43 variants from the GTEx model were available in PAGE summary statistics. In addition to HDL cholesterol levels, *CD36* expression was also associated with levels of C-reactive protein (z-score= 5.30, *P*_TWAS_=1.1×10^-7^).

Although GALA/SAGE multi-ancestry TWAS models showed robust performance, in some cases population-specific models may be preferred to achieve better concordance in ancestry between the training and testing populations. For instance, benign neutropenia is a well-described phenomenon in persons of African ancestry and is almost entirely attributed to variation in the 1q23.2 region. Applying GALA/SAGE AA models to a meta-analysis of 13,476 African Ancestry individuals^31^ identified 139 genes (FDR<0.05), including *ACKR1* (*P*_TWAS_=1.5×10^-234^), the atypical chemokine receptor gene that is the basis of the Duffy blood group system (Figure 5B). This causal gene was missed by GTEx and MESA_AFA_, which detected 100 and 55 genes at FDR<0.05, respectively. TWAS using GALA/SAGE AA also detected 7 genes that were not previously reported in GWAS: *CREB5* (*P*_TWAS_=1.5×10^-14^), *DARS* (*P*_TWAS_=2.9×10^-8^), *CD36* (*P*_TWAS_=1.1×10^-5^), *PPT2* (*P*_TWAS_=1.3×10^-5^), *SSH2* (*P*_TWAS_=4.7×10^-5^), *TOMM5* (*P*_TWAS_=2.9×10^-4^), and *ARF6* (*P*_TWAS_=3.4×10^-4^).

**Figure 5.**
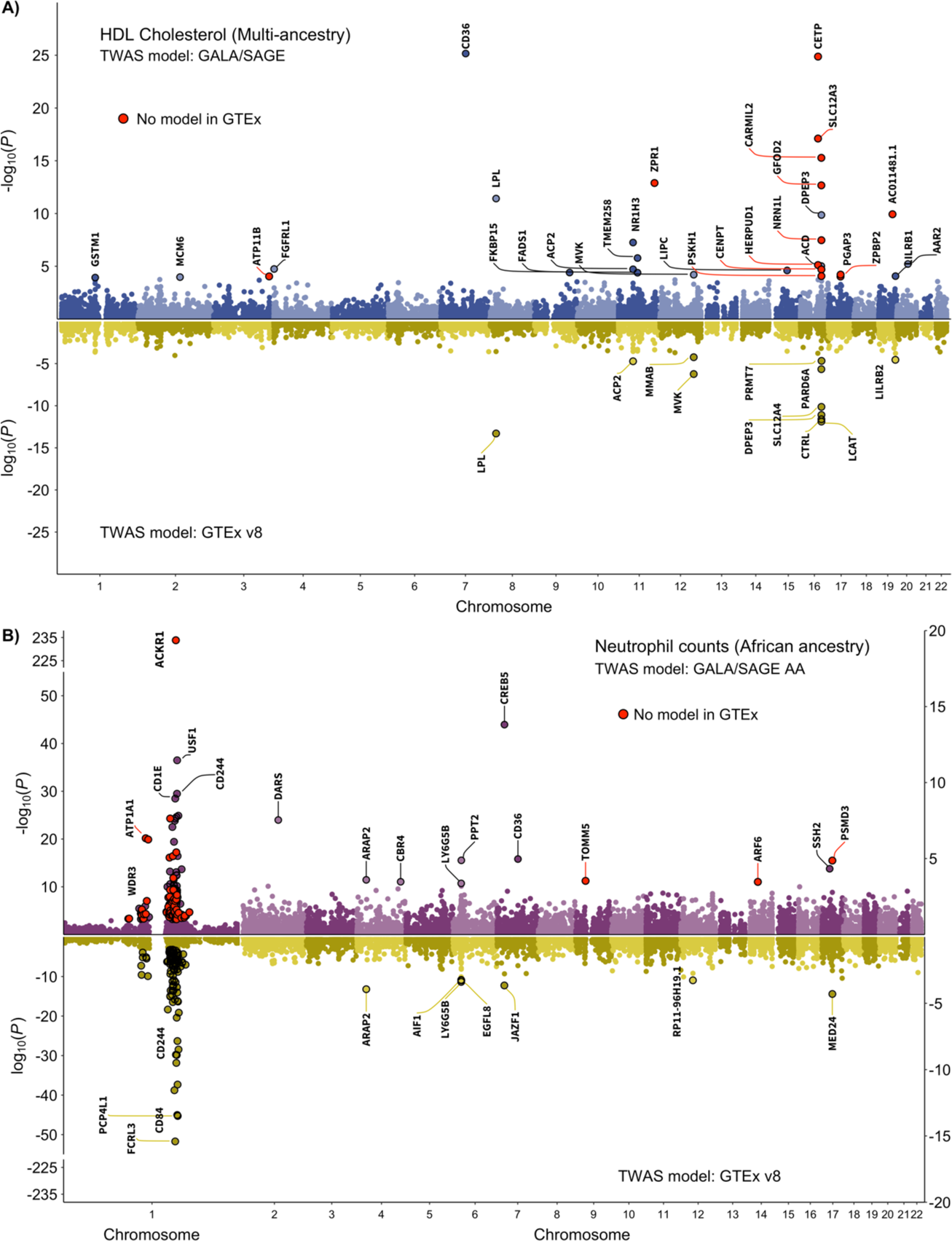
Transcriptome-wide association study (TWAS) results for selected traits. TWAS of HDL in **A)** used GWAS summary statistics from the multi-ancestry PAGE study (N=33,063). TWAS of neutrophil counts in **B)** used summary statistics from a GWAS meta-analysis of African ancestry individuals (N=13,476) by Chen et al. Associated genes (FDR<0.05) are highlighted as circles with a black border and labeled, except for chromosome 1 for neutrophil counts due to the large number of associations. Significantly associated genes for which expression levels could not be predicted using GTEx v8 elastic net models are indicated in red.

Next, we applied GALA/SAGE AA and GTEx models to summary statistics for 22 blood-based biomarkers and quantitative traits from the UK Biobank (UKB). Ancestry-matched TWAS of UKB AFR (median GWAS n=6,190) identified 56 gene-trait associations (FDR<0.05), whereas ancestry-discordant analyses using GTEx detected 92% fewer statistically significant associations, with only 5 genes (Figure S10). TWAS z-scores for associated genes from the two analyses were modesty correlated (*r*=0.37, 95% CI: −0.01 – 0.66). TWAS in UKB EUR (median GWAS n=400,223) also illustrated the advantage of ancestry-matched analyses, but the difference was less dramatic, with a 15% decrease in the number of genes that reached FDR<0.05 using GALA/SAGE AA models, and strong correlation between z-scores (*r*=0.77, 95% CI: 0.76-0.78). With the exception of hemoglobin, where GTEx yielded 1196 genes and AA models detected 326, the number of TWAS-significant findings per trait was comparable. Concordance between significant associations across the 22 traits was 28%, ranging from 1306 (32.7%) genes for height to 108 (7.6%) genes for hemoglobin.

## DISCUSSION

Our comprehensive analysis in a large, multi-racial/multi-ethnic population elucidated the role of genetic ancestry in shaping the genetic architecture of whole blood gene expression that may be applicable to other complex traits. We found that *cis*-heritability of gene expression increased with higher proportion of global African ancestry, and that in admixed populations with intermediate global ancestry, *cis*-heritability was also highest in individuals with predominantly local African ancestry. Parallel analyses of Indigenous American Ancestry revealed an inverse relationship – with genetic variance and *cis*-heritability decreasing in individuals with higher levels of Indigenous American compared to European ancestry. The consistency across analyses of global and local ancestry within self-identified race/ethnicity groups (African Americans or Puerto Ricans) and the pooled GALA/SAGE population suggests that confounding by social or environmental factors is an unlikely explanation for these results. The same pattern was observed for genetic variance, which further supports that differences in heritability between ancestry groups do not simply reflect differences in the relative contribution of environmental factors.

To our knowledge, this relationship between ancestry and heritability has not been previously demonstrated for whole blood gene expression, particularly using WGS data in a sufficiently large and diverse population. Our findings are consistent with the overall pattern of heterozygosity in African and Indigenous American populations. Sub-Saharan African populations consistently show the highest heterozygosity since the ancestors of all other populations passed through a bottleneck during their migration out of Africa^32,33^. Indigenous American populations have passed through additional bottlenecks^34, 35^. With every bottleneck event there is a loss of variation and a concomitant loss of heterozygosity^36^. Therefore, greater genetic control of gene expression in African ancestry populations may be a function of higher heterozygosity resulting in more segregating functional variants in the cis-region^37^. This interpretation is also supported by the higher number of LD-independent *cis*-eQTLs, overall and per-gene, in AFR_high_ compared to AFR_low_ and groups.

A second major finding of our work is that over 30% of heritable protein-coding genes have ancestry-specific eQTLs, most of which are Tier 1 variants that are rare (MAF < 0.01) or even non-polymorphic in another population. The prevalence of the Tier 1 class remained stable when the global ancestry cut-off was increased from 50% to 70% for AFR_high_ and IAM_high_ groups. Our findings align with a recent plasma proteome analysis of the Atherosclerosis Risk in Communities (ARIC) study, which found that nearly 33% of pQTLs identified in a large sample of African Americans (n=1871) were nonexistent or rare in the 1000 Genomes EUR population^38^. Tier 2 anc-eQTLs are an interesting class of variants that are present at a sufficient frequency (MAF>0.01) in both ancestry groups, but do not belong to the same gene-specific credible set. Tier 2 eQTLs could arise due to differences in environmental effects on gene expression, gene-by-gene and gene-by-environment interactions, or multiple causal variants at the same locus that are in different degrees of LD with each other. Among eQTL signals that were shared between ancestry groups effect size heterogeneity was rare. The Tier 3 class of eQTLs was effectively eliminated when AFR_high_ and IAM_high_ were defined using 70% as the global ancestry cut-off, suggesting that heterogeneity in allelic effects is not a major determinant of ancestry-related eQTL differences. However, comparisons of marginal effect sizes are challenging and confounded by differences in sampling error, particularly when there is an imbalance in sample size between populations. Therefore, we may have underestimated ancestry-related heterogeneity in eQTL effects.

Our third major finding relates to the importance of comprehensively accounting for genetic determinants of trait variation in multi-ethnic populations, as illustrated in our TWAS results for 28 traits from the PAGE study. TWAS models trained in the racially/ethnically and ancestrally diverse GALA/SAGE study identified significantly more trait-associated genes than both GTEx and MESA. When applied to admixed populations, GALA/SAGE imputation models benefit from having more similar allele frequency profiles to the target datasets, such as PAGE, as well as more accurate modeling of LD. This is consistent with the findings of Geoffroy et al.^13^ using GTEx and MESA models, as well as other observations^12, 13^ that ancestry-matched models improve power for gene discovery in admixed populations. Over 40% of significantly associated TWAS genes detected using GALA/SAGE models were not available in GTEx, which underscores how biologically meaningful associations may be overlooked in studies that exclusively rely on European ancestry-based predictions. The top two HDL cholesterol-associated genes, *CETP* in 16q13 and *CD36* in 7q21, with established effects on lipid metabolism^19, 39–41^, were not detected in TWAS using GTEx due differences in eQTLs. The finding for *CD36* is compelling since this gene was associated with multiple phenotypes and contains Tier 1 anc-eQTLs that are specific to individuals with >50% African ancestry, consistent with earlier findings that evolutionary pressures have elevated genetic divergence at this locus^42, 43^. *CD36* encodes a transmembrane protein that binds many ligands, including collagen, thrombospondin, and long-chain fatty acids, and also serves as a negative regulator of angiogenesis^44^. Beyond lipid metabolism, the main functions of *CD36* involve mediating the adherence of erythrocytes infected with *Plasmodium falciparum*, the parasite that causes severe malaria^45, 46^.

However, the most striking example of ancestry-specific genetic architecture in our TWAS involves the Duffy antigen receptor gene (*ACKR1*) on 1q23.2, which is responsible for persistently lower white blood cell and neutrophil counts in populations of predominantly African ancestry^47, 48^. Common African-derived alleles at this locus confer a selective advantage against *Plasmodium vivax* malaria and are extremely rare in European ancestry populations. Expression of *ACKR1* could not be imputed using GTEx or MESA, but this causal gene was captured by the pooled and AA-specific GALA/SAGE TWAS models. We also replicated *PSMD3* in 17q21^49^, which was previously identified in African Americans, and several genes that were discovered in European ancestry populations (*CREB5*, *SSH2*, and *PPT2*)^50^. Ancestry-matched TWAS models identified 11 genes associated with neutrophil counts outside of the Duffy locus, including novel genes that have not previously been linked to hematologic traits: *DARS1* in 2q31.1 modulates reactivity to mosquito antigens^51^, while *TOMM5* has been implicated in lipoprotein phospholipase A2 activity^52^.

Our TWAS in UKB illustrated that while ancestry-matched training and testing populations are clearly optimal, there is also evidence that transcriptome prediction models developed in African Americans may have better cross-population portability than models based on predominantly European ancestry samples such as GTEx. Across 22 blood-based biomarkers and traits, the loss of signal was less dramatic in ancestry-discordant analyses that applied models trained in GALA/SAGE African Americans to GWAS summary statistics from UKB EUR subjects than the reverse (15% vs. 92% fewer statistically significant findings). The correlation of TWAS z-scores from ancestry-matched and ancestry-discordant analyses was also lower in UKB AFR than UKB EUR. Similar asymmetric performance has been demonstrated for proteome-wide models in ARIC^37^, where predicted R^2^ standardized by *cis*-h^2^ was higher for AA models applied to EU than for EU modes in AA. We hypothesize that greater genetic diversity of African ancestry populations allows for a more comprehensive set of genetic predictors of transcript levels to be captured by the TWAS models, whereas only a fraction of these variants may be present in populations that underwent additional bottlenecks. Taken together, these findings highlight the value of genetic prediction models trained in ancestrally diverse populations as a resource for identifying trait-associated genes in important biological pathways and advancing research in admixed populations.

PAGE TWAS z-scores were highly correlated across transcriptome models, although the magnitude of correlation with GALA/SAGE was higher for GTEx than MESA_AFHI_ results, which may partly reflect the lack of neutrophils present in monocyte gene expression in MESA_AFHI_ compared to whole blood in GALA/SAGE and GTEx^2^. Furthermore, both GALA/SAGE and GTEx conducted whole-genome sequencing, whereas MESA TWAS models are based on imputed genotype data. However, when comparing GALA/SAGE and MESA there were few instances where a gene was significantly associated based on one model and null using another or associated in both analyses with opposite directions of effect, suggesting that similarity in ancestry may partly compensate for differences in cell type. While our study population is comprised of participants under 21 years of age, TWAS of biomarkers and chronic conditions in adults from PAGE and UKB identified more associated genes than adult-derived prediction models. This implies that the power gained from ancestry-matched models trained in an adequately sized population may outweigh differences in age.

Given that the genetic architecture of complex traits is, to a variable degree, mirrored by the genetics of gene expression^53^, higher heritability in individuals with at least 50% global African ancestry implies that genetic prediction of complex traits should be at least as accurate, if not more effective, in these populations. However, for most complex traits the performance of polygenic prediction models in admixed and predominantly African ancestry individuals lags significantly behind other populations^15^, particularly those of European ancestry, likely due to insufficient sample size and underrepresentation in discovery studies. This is also supported by simulation-based studies and accumulating results from well-powered analyses of diverse cohorts^37, 54, 55^. While these results argue for ancestry-specific estimates of heritability, and the importance of context in heritability estimation, it is important to note that there continues to be a preponderance of relevant ancestry-specific eQTLs across diverse populations. It continues to be important to study and engage with diverse populations across the globe, rather than continue to focus on single-population studies and predictive models.

The substantial prevalence of ancestry-specific eQTLs driven by allele frequency differences also implies that analytic approaches alone will yield limited improvements in the cross-population portability of genetic prediction models, including TWAS and polygenic risk scores. For instance, fine-mapping methods that account for differential LD tagging to identify causal variants will recover some deficits in prediction performance but will not compensate for unobserved risk variants. Our results reinforce the conclusion that developing truly generalizable genetic prediction models requires capturing the full spectrum of genetic variation across human populations. As such, access to sufficiently large ancestrally diverse populations remains the main rate-limiting step.

In evaluating the contributions of our work, several limitations should be acknowledged. Our study was limited to whole blood and similar analyses of ancestry-specific effects should be performed for other tissues. However, whole blood is one of the most clinically-informative and commonly-collected samples, and for over 60% of genes whole blood transcriptomes significantly capture expression levels in other tissues^56^. Thus, our observations regarding the genetic architecture of whole blood eQTLs in admixed populations with African and Indigenous American ancestry are likely generalizable to other tissues. Our approach for classifying ancestry-specific eQTLs may result in an underestimation of the number of these loci. We assumed that each gene had one causal eQTL locus and focused all comparisons on the corresponding 95% credible set. This assumption is likely violated for genes with multiple independent eQTLs, which would limit our ability to assess the ancestry-specificity of all signals. We believe this is a conservative assumption that would lead us to potentially miss some ancestry-specific eQTLs. Detection of our Tier 2 anc-eQTLs by PESCA relies on having regions that are approximately LD independent in both populations to estimate the proportion of causal variants. This estimate may be biased if there is residual LD between regions, which is a challenge in admixed populations with longer-range LD. Lastly, our comparison of TWAS models may be slightly biased against GTEx in European ancestry TWAS since we did not apply MASHR models, which predict a larger number of genes using fine-mapped eQTLs^57^. We chose to compare with elastic net GTEx models because GALA/SAGE TWAS models were developed using the same analytic pipeline.

Although there is evidence that accounting for local ancestry increases power for discovery in cis-eQTL mapping^58, 59^, adjustment for local ancestry as a covariate did not improve the predictive performance of TWAS models. Previous work by Gay *et al.* reported that local ancestry explains at least 7% of the variance in residual expression for 1% of expressed genes in 117 admixed individuals from GTEx ^58^. In GALA II/SAGE, we found that local ancestry was a significant predictor of transcript levels for at least 10% of heritable genes, explaining between 2.1% (in 893 Puerto Ricans) and 5.1% (in 757 African Americans) of residual variance. Consistent with Gay et al., we observed that local ancestry explains a larger proportion of variance in gene expression corrected for global ancestry. However, it is possible that the lack of improvement in the TWAS context may be due to overadjustment as local ancestry may serve as a proxy for information already captured by population-specific genetic variants, or because of how local ancestry was modelled in our analyses.

Despite these limitations, our study leveraged a uniquely large and diverse sample of 2,733 African American and Latino participants to explore the interplay between genetic ancestry and regulation of gene expression. Our approach to evaluating the degree of specificity of whole blood eQTLs to African or Indigenous American ancestry revealed that such effects are mostly driven by allele frequency differences between populations. Tier 1 anc-eQTLs reach a frequency of at least 1% only in predominantly African or Indigenous American ancestry populations and affect the expression of a large fraction of protein-coding genes, which has implications for detecting functional genetic variants and evaluating their role in disease susceptibility. In addition, we provide genetic prediction models of whole blood transcriptomes that cover a greater number of genes than similar resources developed in European ancestry populations and facilitate more powerful TWAS when applied to studies of admixed individuals and multi-ancestry GWAS meta-analyses. In summary, our study highlights the need for larger genomic studies in globally representative populations for characterizing the genetic basis of complex traits and ensuring equitable translation of precision medicine efforts.

## METHODS

### Study population

This study examined African American, Puerto Rican and Mexican American children between 8-21 years of age with or without physician-diagnosed asthma from the Genes-environments and Admixture in Latino Americans II (GALA II) study and the Study of African Americans, Asthma, Genes & Environments (SAGE). The inclusion and exclusion criteria are previously described in detail^60, 61^. Briefly, participants were eligible if they were 8-21 years of age and identified all four grandparents as Latino for GALA II or African American for SAGE. Study exclusion criteria included the following: 1) any smoking within one year of the recruitment date; 2) 10 or more pack-years of smoking; 3) pregnancy in the third trimester; 4) history of lung diseases other than asthma (for cases) or chronic illness (for cases and controls). The local institutional review board from the University of California San Francisco Human Research Protection Program approved the studies (IRB# 10-02877 for SAGE and 10-00889 for GALA II). All subjects and their legal guardians provided written informed consent.

### Whole genome sequencing data and processing

Genomic DNA samples extracted from whole blood were sequenced as part of the Trans-Omics for Precision Medicine (TOPMed) whole genome sequencing (WGS) program^62^ and the Centers for Common Disease Genomes of the Genome Sequencing Program. WGS was performed at the New York Genome Center and Northwest Genomics Center on a HiSeq X system (Illumina, San Diego, CA) using a paired-end read length of 150 base pairs (bp), with a minimum of 30x mean genome coverage. DNA sample handling, quality control, library construction, clustering, and sequencing, read processing and sequence data quality control are previously described in detail^62^. All samples were jointly genotyped by the TOPMed Informatics Research Center. Variant calls were obtained from TOPMed data freeze 8 VCF files generated based on the GRCh38 assembly. Variants with a minimum read depth of 10 (DP10) were used for analysis unless otherwise stated.

### RNA sequencing data generation and processing

Total RNA was isolated from PAXgene tube using MagMax^TM^ for Stabilized Blood Tubes RNA Isolation Kit (Applied Biosystem, P/N 4452306). Globin depletion was performed using GLOBINcleasr^TM^ Human (Thermo Fisher Scientific, cat. no. AM1980). RNA integrity and yield were assessed using an Agilent 2100 Bioanalyzer (Agilent Technologies, Santa Clara, CA, USA).

Total RNA was quantified using the Quant-iT™ RiboGreen® RNA Assay Kit and normalized to 5ng/ul. An aliquot of 300ng for each sample was transferred into library preparation which was an automated variant of the Illumina TruSeq™ Stranded mRNA Sample Preparation Kit. This method preserves strand orientation of the RNA transcript. It uses oligo dT beads to select mRNA from the total RNA sample. It is followed by heat fragmentation and cDNA synthesis from the RNA template. The resultant cDNA then goes through library preparation (end repair, base ‘A’ addition, adapter ligation, and enrichment) using Broad-designed indexed adapters substituted in for multiplexing. After enrichment the libraries were quantified with qPCR using the KAPA Library Quantification Kit for Illumina Sequencing Platforms and then pooled equimolarly. The entire process is in 96-well format and all pipetting is done by either Agilent Bravo or Hamilton Starlet.

Pooled libraries were normalized to 2nM and denatured using 0.1 N NaOH prior to sequencing. Flowcell cluster amplification and sequencing were performed according to the manufacturer’s protocols using the HiSeq 4000. Each run was a 101bp paired-end with an eight-base index barcode read. Each sample was targeted to 50M reads. Data was analyzed using the Broad Picard Pipeline which includes de-multiplexing and data aggregation.

RNA-seq reads were further processed using the TOPMed RNA-seq pipeline for Year 3 and Phase 5 RNA-seq data (supplementary file 2 obtained from https://topmed.nhlbi.nih.gov/sites/default/ files/TOPMed_RNAseq_pipeline_COREyr3.pdf). Count-level data were generated using GRCh38 human reference genome and GENCODE 30 for transcript annotation. Count-level quality control (QC) and normalization were performed following the Genotype-Tissue Expression (GTEx) project v8 protocol (https://gtexportal.org/home/methods). Sample-level QC included removal of RNA samples with RIN < 6, genetically related samples (equal or more related than third degree relative), and sex-discordant samples based on reported sex and their *XIST* and *RPS4Y1* gene expression profiles. Count distribution outliers were detected as follows: (i) Raw counts were normalized using the trimmed mean of M values (TMM) method in edgeR^63^ as described in GTEx v8 protocol. (ii) The log2 transformed normalized counts at the 25th percentile of every sample were identified (count_q25_). (iii) The 25th percentile (Q25) of count_q25_ was calculated. (iv) Samples were removed if their count_q25_ was lower than −4 as defined by visual inspection.

To account for hidden confounding factors such as batch effects, technical and biological variation in the sample preparation, and sequencing and/or data processing procedures, latent factors were estimated using the Probabilistic Estimation of Expression Residuals (PEER) method^64^. Optimization was performed according to approach adopted by GTEx with the goal to maximized eQTL discovery^65^. A total of 50 (for AA, PR, MX, pooled samples) and 60 (for AFR_high_, AFR_low_, IAM_high_, IAM_low_) PEER factors were selected for downstream analyses (Figure S11).

### Estimation of global and local genetic ancestry

Genetic principal components (PCs), global and local ancestry, and kinship estimation on genetic relatedness were computed using biallelic single nucleotide polymorphisms (SNPs) with a PASS flag from TOPMed freeze 8 DP10 data as described previously^66, 67^. Briefly, genotype data from European, African, and Indigenous American (IAM) ancestral populations were used as the reference panels for global and local ancestry estimation assuming three ancestral populations.

Reference genotypes for European (HapMap CEU) and African (HapMap YRI) ancestries were obtained from the Axiom® Genotype Data Set (https://www.thermofisher.com/us/en/home/life-science/microarray-analysis/microarray-data-analysis/microarray-analysis-sample-data/axiom-genotype-data-set.). The CEU populations were recruited from Utah residents with Northern and Western European ancestry from the CEPH collection. The YRI populations were recruited from Yoruba in Ibadan, Nigeria. The Axiom® Genome-Wide LAT 1 array was used to generate the Indigenous American (IAM) ancestry reference genotypes from 71 Indigenous Americans (14 Zapotec, 2 Mixe and 11 Mixtec from Oaxaca, 44 Nahua from Central Mexico)^68, 69^. ADMIXTURE was used with the reference genotypes in a supervised analysis assuming three ancestral populations. Global ancestry was estimated by ADMIXTURE^70^ in supervised while local ancestry was estimated by RFMIX version 2 with default settings^71^. Throughout this study, local ancestry of a gene was defined as the number of ancestral alleles (0, 1, or 2) at the transcription start site.

Comparative analyses were performed based on two different sample grouping strategies, by self-identified race/ethnicity or by global ancestry. Self-identified race/ethnicity included four groups – African Americans (AA), Puerto Ricans (PR), Mexican Americans (MX), and the pooling of AA, PR, MX and other Latinos (pooled). For groups defined by global ancestry, samples were grouped into high (> 50%, AFR_high_ or IAM_high_) or low (< 10%, AFR_low_ or IAM_low_) global African or Indigenous American ancestry. The sample size for each group is shown in Table S1.

### Cis-heritability of gene expression

The genetic region of *cis*-gene regulation was defined by 1MB region flanking each side of the transcription start site (*cis*-region). *Cis*-heritability (h^2^) of gene expression was estimated using unconstrained GREML^72^ analysis (--reml-no-constrain), and estimation was restricted to common autosomal variants (MAF ≥ 0.01). Inverse-normalized gene expression was regressed on PEER factors, and the residuals were used as the phenotype for GREML analysis. Sex and asthma case-control status was used as categorical covariates, while age at blood draw and the first 5 genetic PCs were used as quantitative covariates. *Cis*-heritability was estimated separately for each self-identified race/ethnicity group (AA, PR, MX and pooled) and groupings based on global (AFR_high_, AFR_low_, IAM_high_ and IAM_low_) and local ancestry (described below). Differences in the distribution of h^2^ and genetic variance (V_G_) between groups were tested using two-sided Wilcoxon tests. Parallel analyses were also conducted for Indigenous American ancestry (IAM/IAM vs. EUR/EUR and IAM/IAM vs. IAM/EUR).

The following sensitivity analyses were conducted using GCTA: i) using the same sample size in each self-identified group (n=600) and (ii) partitioning heritability and genetic variance by two minor allele frequency bins (0.01-0.1, 0.1-0.5). We also estimated heritability using the LDAK-Thin model^73^, following the recommended GRM processing. Thinning of duplicate SNPs was performed using the arguments “--window-prune .98 -- window-kb 100”. The direct method was applied to calculate kinship using the thinned data and lastly, generalized restricted maximum likelihood (REML) was used to estimate heritability.

### Association of global and local ancestry with gene expression

Methods from Gay *et al* (2020)^58^ was modified to identify genes associated with global and local ancestry (Figure S1). In step 1, inversed normalized gene expression was regressed on age, sex and asthma status (model 0). In step 2, the residuals from model 0 were regressed on global ancestry (model 1). In step 3, the residuals from model 1 were regressed on local ancestry (model 2) to identify genes that are associated with local ancestry. A false discovery rate (FDR) of 0.05 was applied to step 2 and 3 separately to identify genes that were significantly associated with global and/or local ancestry. Step 1 to step 3 were run separately for African and Indigenous American ancestry. For heritable genes that were associated with global and/or local ancestry, a joint model of regressing global and local ancestry from residuals from model 0 was also examined to assess the percentage of variance of gene expression explained by global and/or local ancestry.

### Identification of eGenes, cis-eQTLs and ancestry-specific cis-eQTLs

Raw gene counts were processed and eQTLs were identified using FastQTL^74^ according to the GTEx v8 pipeline (https://github.com/broadinstitute/gtex-pipeline). Age, sex, asthma status, first 5 genetic ancestry PCs, and PEER factors were used as covariates for FastQTL analysis. To account for multiple testing across all tested genes, the Benjamini & Hochberg correction was applied to the beta-approximated p-values from the permutation step of FastQTL. For each gene with a significant beta-approximated p-value at the false discovery rate < 0.05, a nominal p-value threshold was estimated using the beta-approximated p-value. *Cis*-eQTLs were defined as genetic variants that have nominal p-values less than the nominal p-value threshold of the corresponding gene. eGenes were defined as genes with at least one eQTL. To summarize the number of independent cis-eQTLs in each ancestry group, LD clumping was performed using PLINK (--clump-kb 1000 -- clump-r2 0.1) using gene-specific p-value thresholds.

*Trans*-eQTLs were identified using the same protocol as in GTEx v8^2^. *Trans*-eQTLs were defined as eQTLs that were not located on the same chromosome as the gene. Only protein-coding and lincRNA genes and SNPs on autosomes were included in the analyses. Briefly, linear regression on expression of gene was performed in PLINK2 (version v2.00a3LM released 28 Mar 2020) using SNPs with MAF ≥ 0.05 and the same covariates as *cis*-eQTL discovery. Gene and variant mappability data (GRCh38 and GENCODE v26) were downloaded from Saha and Battle^75^ for the following filtering steps: (i) keep gene-variant pairs that passed a p–value threshold of 1×10^-5^, (ii) keep genes with mappability ≥ 0.8, (iii) remove SNPs with mappability < 1, and (iv) remove a *trans*-eQTL candidate if genes within 1MB of the SNP candidate cross-mapped with the trans-eGene candidate. The Benjamini-Hochberg procedure was applied to control for FDR at the 0.05 level using the smallest p-value (multiplied by 10^-6^) from each gene. An additional filtering step was applied for the AFR_high_ and IAM_high_ groups. For AFR_high_, all trans-eQTLs detected in AFR_low_ were removed and the resulting trans-eQTL were referred to as filtered AFR_high_ trans-eQTLs. Similarity, for IAM_high_ groups, all trans-eQTLs detected in IAM_low_ groups were removed and the resulting trans-eQTL were referred to as filtered IAM_high_ trans-eQTLs. Filtered AFR_high_ trans-eQTL were checked for presence of filtered IAM_high_ trans-eQTLs, and vice versa. LD clumping was performed using PLINK (v1.90b6.26 --clump-kb 1000 --clump-r2 0.1 --clump-p1 0.00000005 -- clump-p2 1) to group trans-eQTLs into independent signals.

Ancestry-specific eQTL (anc-eQTL) mapping was performed in participants stratified by high and low global African and Indigenous American ancestry (see “Grouping samples by self-identified race/ethnicity or global ancestry”). We developed a framework to identify anc-eQTLs by focusing on the lead eQTL signal for each gene and comparing fine-mapped 95% credible sets between high (>50%) and low (<10%) global ancestry groups (AFR_high_ vs AFR_low_; IAM_high_ vs IAM_low_). Sensitivity analyses were conducted using >70% as the cut-off for AFR_high_ and IAM_high_ groups. Anc-eQTLs were classified into three tiers as described below, based on population differences in allele frequency, linkage disequilibrium (LD), and effect size (Figure 3A). For every protein-coding and heritable eGene (GCTA h^2^ LRT p-value <0.05), the lead eQTL signal was identified using CAVIAR^76^ assuming one causal locus (c=1). The 95% credible set of eQTLs in the high and low global ancestry group were compared to determine if there was any overlap. Variants from non-overlapping 95% credible sets were further classified as Tier 1 anc-eQTLs based on allele frequency differences or Tier 2 after additional fine-mapping using PESCA^21^. For genes with overlapping 95% credible sets, Tier 3 anc-eQTLs were detected based on effect size heterogeneity.

eQTLs identified in AFR_high_ or IAM_high_ high group that were common (MAF ≥0.01) in the high group but rare (MAF<0.01) or monomorphic in the AFR_low_ or IAM_low_ group were classified as Tier 1. If the eQTLs were detected at MAF≥0.01 in both the high and low ancestry groups, they were further fine-mapped using PESCA^21^, which tests for differential effect sizes while accounting for LD between eQTLs. Pre-processing for the PESCA analyses involved LD pruning at r2 >0.95. All eQTL pairs with r2 >0.95 were identified in both the high and low groups and only those pairs common to both groups were removed. For each eQTL, PESCA estimated three posterior probabilities: specific to the AFR_high_ or IAM_high_ group (PP_high_), specific to the AFR_low_ or IAM_low_ group (PP_low_), or shared between the two groups (PP_shared_). Tier 2 anc-eQTLs were selected based on the following criteria: i) all variants in the credible set had (PP_high_ > PP_low_) and (PP_high_ > PP_shared_) and ii) PP_high_ > 0.8. Tier 3 class was based on evidence of significant heterogeneity in eQTL effect size, defined as Cochran’s Q p-value < 0.05/nGene, where nGene was the number of genes tested. Since we assume the 95% credible set corresponds to a single lead eQTL signal, all eQTLs in the credible set were required to have a significant heterogeneous effect size to be classified as Tier 3 anc-eQTLs.

To systematically assess the overlap in eQTL signals identified in our study and trait-associated loci, we colocalized eQTL summary statistics with GWAS results from PAGE. Colocalization was performed using COLOC^77^ within a LD window of 2 MB centered on the eQTL with the lowest GWAS p-value. For each eQTL-trait pair, the posterior probably of a shared causal signal (PP_4_) >0.80 was interpreted as strong evidence of colocalization.

### Development of gene prediction models and transcriptome-wide association analyses

Gene prediction models for *cis*-gene expression were generated using common variants and elastic net modeling implemented in the PredictDB v7 pipeline (https://github.com/hakyimlab/ PredictDB_Pipeline_GTEx_v7). Models were filtered by nested cross validation (CV) prediction performance and heritability p-value (rho_avg > 0.1, zscore_pval <0.05 and GCTA h^2^ p-value < 0.05). Sensitivity analyses were performed by generating gene prediction models that included the number of ancestral alleles as covariates to account for local ancestry in the *cis*-region. In AA, one covariate indicating the count of African ancestral allele was used while in PR, MX, and pooled, two additional covariates indicating the number of European and Indigenous American ancestral alleles were used.

Out-of-sample validation of the gene expression prediction models were done using 598 individuals from the African American asthma cohort, Study of Asthma Phenotypes and Pharmacogenomic Interactions by Race-Ethnicity (SAPPHIRE)^30^. Predicted gene expression from SAPPHIRE genotypes was generated using the predict function from MetaXcan. Genotypes of SAPPHIRE samples were generated by whole genome sequencing through the TOPMed program and were processed the same way as GALA II and SAGE. RNA-seq data from SAPPHIRE were generated as previously described^78^ and were normalized using TMM in edgeR. Predicted and normalized gene expression data were compared to generate correlation R^2^.

To assess the performance of the resulting GALA/SAGE models we conducted transcriptome-wide association studies (TWAS) of 28 traits using GWAS summary statistics from the Population Architecture using Genomics and Epidemiology (PAGE) Consortium study by Wojcik et al^20^. Analyses were performed using S-PrediXcan with whole blood gene prediction models from GALA II and SAGE (GALA/SAGE models), GTEx v8, and monocyte gene expression models from the Multi-Ethnic Study of Atherosclerosis (MESA) study^3^. In the UK Biobank we conducted TWAS of 22 blood-based biomarkers and quantitative traits using GALA/SAGE models generated in African Americans (GALA/SAGE AA) and GTEx v8 whole blood. Each set of TWAS models was applied to publicly available GWAS summary statistics (Pan-UKB team: https://pan.ukbb.broadinstitute.org) from participants of predominantly European ancestry (UKB EUR) and African ancestry (UKB AFR). Ancestry assignment in UKB was based on a random forest classifier trained on the merged 1000 Genomes and Human Genome Diversity Project (HGDP) reference populations. The classifier was applied to UK Biobank participants projected into the 1000G and HGDP principal components.

## Data availability

TOPMed WGS and RNA-seq data from GALA II and SAGE are available on dbGaP under accession number phs000920.v4.p2 and phs000921.v4.p1, respectively. TOPMed WGS data from SAPPHIRE are available under the dbGaP accession number phs001467.v1.p1. Summary statistics for cis- and trans-eQTLs, as well as TWAS models developed using data from GALA II and SAGE participants have been posted in the following public repository DOI: 10.5281/zenodo.6622368

## ACKNOWLEDGEMENTS

Generation of molecular data for the TOPMed (Trans-Omics in Precision Medicine) program was supported by the National Heart, Lung and Blood Institute (NHLBI). RNA sequencing for “NHLBI TOPMed: Gene-Environment, Admixture and Latino Asthmatics Study” (phs000920, GALA II) and “NHLBI TOPMed: Study of African Americans, Asthma, Genes and Environments” (phs000921, SAGE) was performed at Broad Institute Genomics Platform (HHSN268201600034I). Whole genome sequencing (WGS) for the same studies were performed at the New York Genome Center (NYGC, 3R01HL117004-02S3) and Northwest Genomics Center (NWGC, HHSN268201600032I). Core support including centralized genomic read mapping and genotype calling, along with variant quality metrics and filtering were provided by the TOPMed Informatics Research Center (3R01HL-117626-02S1; contract HHSN268201800002I). Core support including phenotype harmonization, data management, sample-identity QC, and general program coordination were provided by the TOPMed Data Coordinating Center (R01HL-120393; U01HL-120393; contract HHSN268201800001I).

Whole genome sequencing of part of GALA II was performed by the New York Genome Center under The Centers for Common Disease Genomics of the Genome Sequencing Program (GSP) Grant (UM1 HG008901). The GSP Coordinating Center (U24 HG008956) contributed to cross-program scientific initiatives and provided logistical and general study coordination. GSP is funded by the National Human Genome Research Institute, the National Heart, Lung, and Blood Institute, and the National Eye Institute.

This work and EGB were supported in part by the Sandler Family Foundation, the American Asthma Foundation, the RWJF Amos Medical Faculty Development Program, Harry Wm. and Diana V. Hind Distinguished Professor in Pharmaceutical Sciences II, the National Heart, Lung, and Blood Institute (R01HL117004, R01HL135156, X01HL134589, U01HL138626), the National Institute of Health and Environmental Health Sciences (R01ES015794), the National Institute on Minority Health and Health Disparities (R56MD013312, P60MD006902), the Tobacco-Related Disease Research Program (24RT-0025, 27IR-0030), and the National Human Genome Research Institute (U01HG009080). LK was supported by funding from National Cancer Institute (K99CA246076). KLK was additionally supported by a diversity supplement of NHLBI R01HL135156, the UCSF Bakar Computational Health Sciences Institute, the Gordon and Betty Moore Foundation grant GBMF3834, and the Alfred P. Sloan Foundation grant 2013-10-27 to UC Berkeley through the Moore-Sloan Data Sciences Environment initiative at the Berkeley Institute for Data Science (BIDS). CRG was supported by funding from the National Human Genome Research Institute (R01HG010297), CRG and NAZ were supported by funding from the National Heart, Lung, and Blood Institute (R01HL151152) and the National Human Genome Research Institute (R01HG011345). We gratefully acknowledge the studies and participants who provided biological samples and data for TOPMed. We thank Soren Germer, Michael C. Zody, Lara Winterkorn and Catherine Reeves from NYGC and Deborah A. Nickerson from NWGC for overseeing the production of GALA II and SAGE whole genome sequencing data. We thank Huwenbo Shi, Kathryn S. Burch and Bodgdan Pasaniuc for their technical support on the PESCA software. The content is solely the responsibility of the authors and does not necessarily represent the official views of the National Institutes of Health

## AUTHOR CONTRIBUTIONS

ACYM, LK, KLK, CRG, NZ, EGB, EZ contributed to the conception or design of the work. LK, ACYM, DH, CE, SH, JRE, NG, SG, SX, HG, AOO, JRS, MAL, LKW, LNB, CRG, NZ, EGB, EZ contributed to the acquisition, analysis, or interpretation of data. LK, ACYM, JRE, KLK, AOO, LNB, CRG, NZ, EGB, EZ have drafted the work or substantively revised it. All authors approved the submission of this manuscript.

---

**Table S1:**
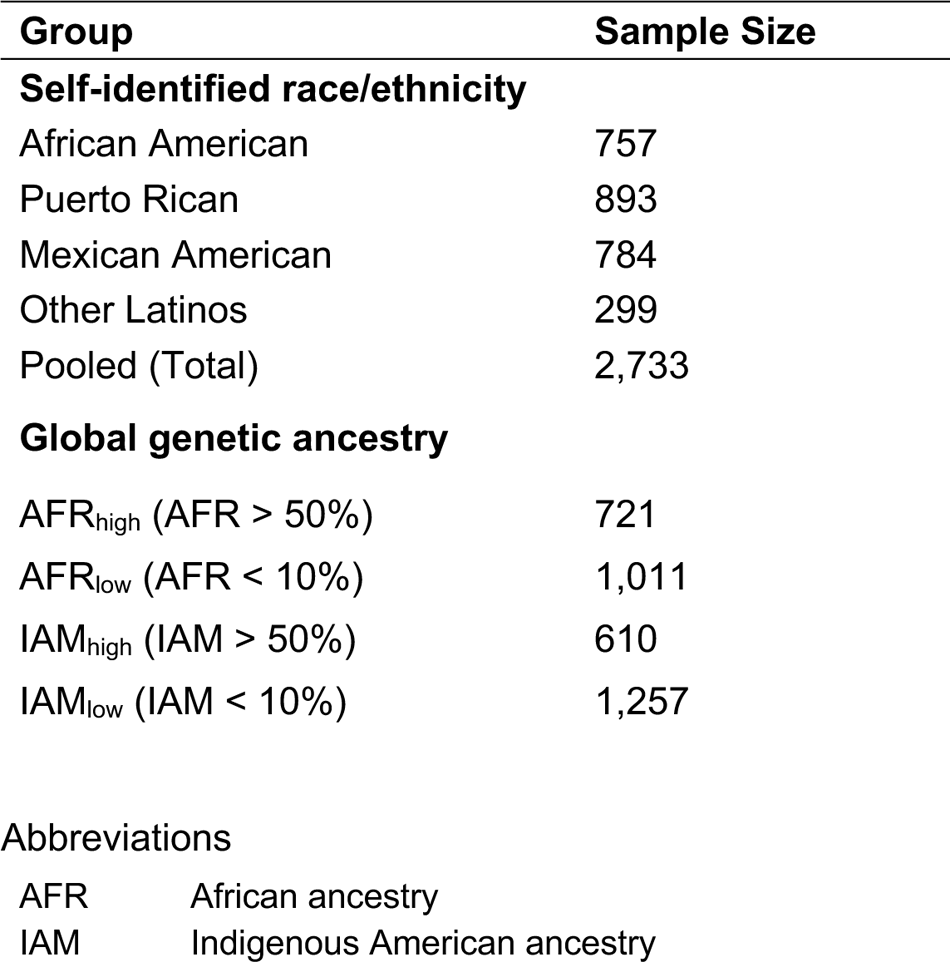
Sample size overview. The total number of individuals with WGS and RNA-seq data that were included in analyses based on self-identified race/ethnicity and genetic ancestry.

| Group | Sample Size |
| --- | --- |
| **Self-identified race/ethnicity** |  |
| African American | 757 |
| Puerto Rican | 893 |
| Mexican American | 784 |
| Other Latinos | 299 |
| Pooled (Total) | 2,733 |
| **Global genetic ancestry** |  |
| AFR<sub>high</sub> (AFR > 50%) | 721 |
| AFR<sub>low</sub> (AFR < 10%) | 1,011 |
| IAM<sub>high</sub> (IAM > 50%) | 610 |
| IAM<sub>low</sub> (IAM < 10%) | 1,257 |
| AFR | African ancestry |
| IAM | Indigenous American ancestry |

**Table S2:**
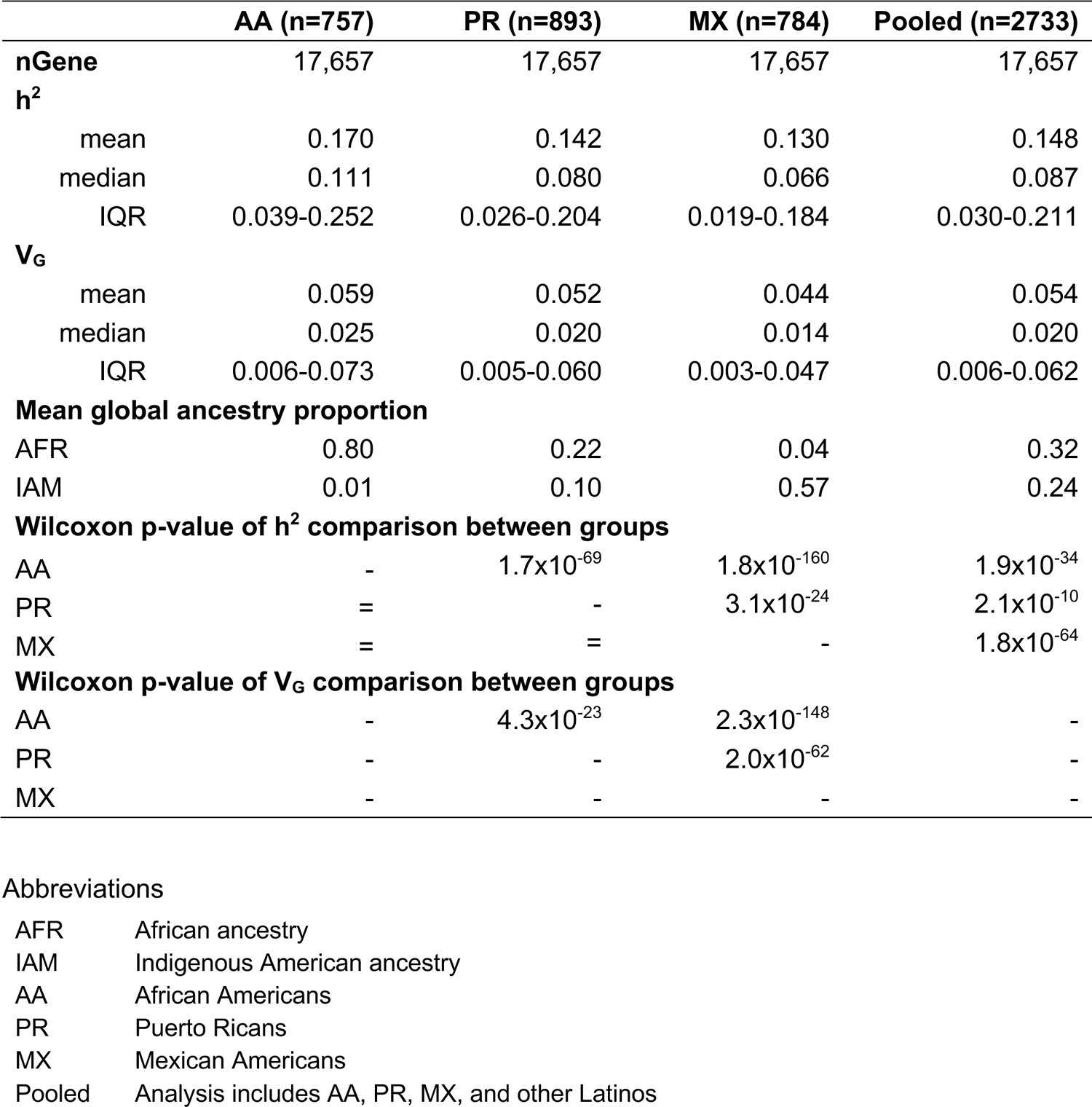
*Cis*-heritability (h^2^) and genetic variance (V_G_) of gene expression stratified by self-identified race/ethnicity. GCTA analyses were restricted to common variants (MAF >= 0.01) in each population within 1MB flanking regions of the transcription start site. Estimates of h^2^ and V_G_ are summarized across the intersection of genes (nGene) with GCTA results available in all populations.

|  | AA (n=757) | PR (n=893) | MX (n=784) | Pooled (n=2733) |
| --- | --- | --- | --- | --- |
| **nGene** | 17,657 | 17,657 | 17,657 | 17,657 |
| **$h^2$** |  |  |  |  |
| mean | 0.170 | 0.142 | 0.130 | 0.148 |
| median | 0.111 | 0.080 | 0.066 | 0.087 |
| IQR | 0.039-0.252 | 0.026-0.204 | 0.019-0.184 | 0.030-0.211 |
| **$V_G$** |  |  |  |  |
| mean | 0.059 | 0.052 | 0.044 | 0.054 |
| median | 0.025 | 0.020 | 0.014 | 0.020 |
| IQR | 0.006-0.073 | 0.005-0.060 | 0.003-0.047 | 0.006-0.062 |
| **Mean global ancestry proportion** |  |  |  |  |
| AFR | 0.80 | 0.22 | 0.04 | 0.32 |
| IAM | 0.01 | 0.10 | 0.57 | 0.24 |
| **Wilcoxon p-value of $h^2$ comparison between groups** |  |  |  |  |
| AA | - | $1.7 \times 10^{-69}$ | $1.8 \times 10^{-160}$ | $1.9 \times 10^{-34}$ |
| PR | = | - | $3.1 \times 10^{-24}$ | $2.1 \times 10^{-10}$ |
| MX | = | = | - | $1.8 \times 10^{-64}$ |
| **Wilcoxon p-value of $V_G$ comparison between groups** |  |  |  |  |
| AA | - | $4.3 \times 10^{-23}$ | $2.3 \times 10^{-148}$ | - |
| PR | - | - | $2.0 \times 10^{-62}$ | - |
| MX | - | - | - | - |

|  |  |
| --- | --- |
| AFR | African ancestry |
| IAM | Indigenous American ancestry |
| AA | African Americans |
| PR | Puerto Ricans |
| MX | Mexican Americans |
| Pooled | Analysis includes AA, PR, MX, and other Latinos |

**Table S3:** *Cis*-heritability (h^2^) and genetic variance (V_G_) of gene expression stratified by global genetic ancestry. GCTA analyses were restricted to common variants (MAF >= 0.01) in each population within 1MB flanking regions of the transcription start site. Individuals were stratified based on proportion. Individuals with >50% global genetic African ancestry (AFR_high_) were compared to those with <10% (AFR_low_). Individuals with >50% global genetic Indigenous American ancestry (IAM_high_) were compared to those with <10% (IAM_low_). Estimates of h^2^ and V^G^ are summarized across the intersection of genes (nGene) with GCTA results available in all genetic ancestry groups.

|  | **$AFR_{high}$ (n=721)** | **$AFR_{low}$ (n=1011)** | **$IAM_{high}$ (n=610)** | **$IAM_{low}$ (n=1257)** |
| --- | --- | --- | --- | --- |
| **nGene** | 18,725 | 18,725 | 18,725 | 18,725 |
| **$h^2$** |  |  |  |  |
| mean | 0.167 | 0.129 | 0.123 | 0.152 |
| median | 0.107 | 0.065 | 0.062 | 0.091 |
| IQR | 0.037-0.247 | 0.019-0.182 | 0.016-0.176 | 0.031-0.221 |
| **$V_G$** |  |  |  |  |
| mean | 0.058 | 0.046 | 0.041 | 0.055 |
| median | 0.024 | 0.014 | 0.013 | 0.022 |
| IQR | 0.006-0.072 | 0.003-0.048 | 0.002-0.045 | 0.006-0.066 |
| **Mean global ancestry proportion** |  |  |  |  |
| AFR | 0.82 | 0.04 | 0.04 | 0.58 |
| IAM | 0.01 | 0.54 | 0.67 | 0.04 |
| **Wilcoxon p-value of $h^2$ comparison between groups** |  |  |  |  |
| $AFR_{high}$ | - | $4.2 \times 10^{-140}$ | - | - |
| $IAM_{high}$ | - | - | - | $5.8 \times 10^{-120}$ |
| **Wilcoxon p-value of $V_G$ comparison between groups** |  |  |  |  |
| $AFR_{high}$ | - | $8.0 \times 10^{-114}$ | - | - |
| $IAM_{high}$ | - | - | - | $8.3 \times 10^{-173}$ |

**Table S4:** Comparison of V_G_ stratified by local genetic ancestry. GCTA analyses were restricted to common variants (MAF >= 0.01) in each population within 1MB flanking regions of the transcription start site. For each gene, individuals were classified into local ancestry groups, L1 and L2, based on the ancestry at the transcription start site. The number of genes (nGene) for which GCTA models successfully converged and produced reliable estimates is reported for each analysis. Genes were not filtered based on heritability.

| Group | Local ancestry | | Mean $h^2$ | | Mean $V_G$ | | nGene | Sample size | Wilcoxon p-value |
| --- | --- | --- | --- | --- | --- | --- | --- | --- | --- |
|  | L1 | L2 | L1 | L2 | L1 | L2 |  |  |  |
| Pooled | AFR/AFR | AFR/EUR | 0.153 | 0.142 | 0.053 | 0.049 | 17,866 | 516 | $2.0 \times 10^{-7}$ |
| AA | AFR/AFR | AFR/EUR | 0.143 | 0.137 | 0.041 | 0.039 | 19,224 | 202 | $5.7 \times 10^{-5}$ |
| PR | AFR/EUR | EUR/EUR | 0.129 | 0.108 | 0.041 | 0.033 | 18,570 | 242 | $1.4 \times 10^{-28}$ |
| Pooled | IAM/IAM | IAM/EUR | 0.108 | 0.119 | 0.033 | 0.038 | 10,566 | 359 | $1.6 \times 10^{-8}$ |
| MX | IAM/IAM | IAM/EUR | 0.101 | 0.117 | 0.029 | 0.035 | 18,194 | 262 | $7.7 \times 10^{-11}$ |

**Table S5:**
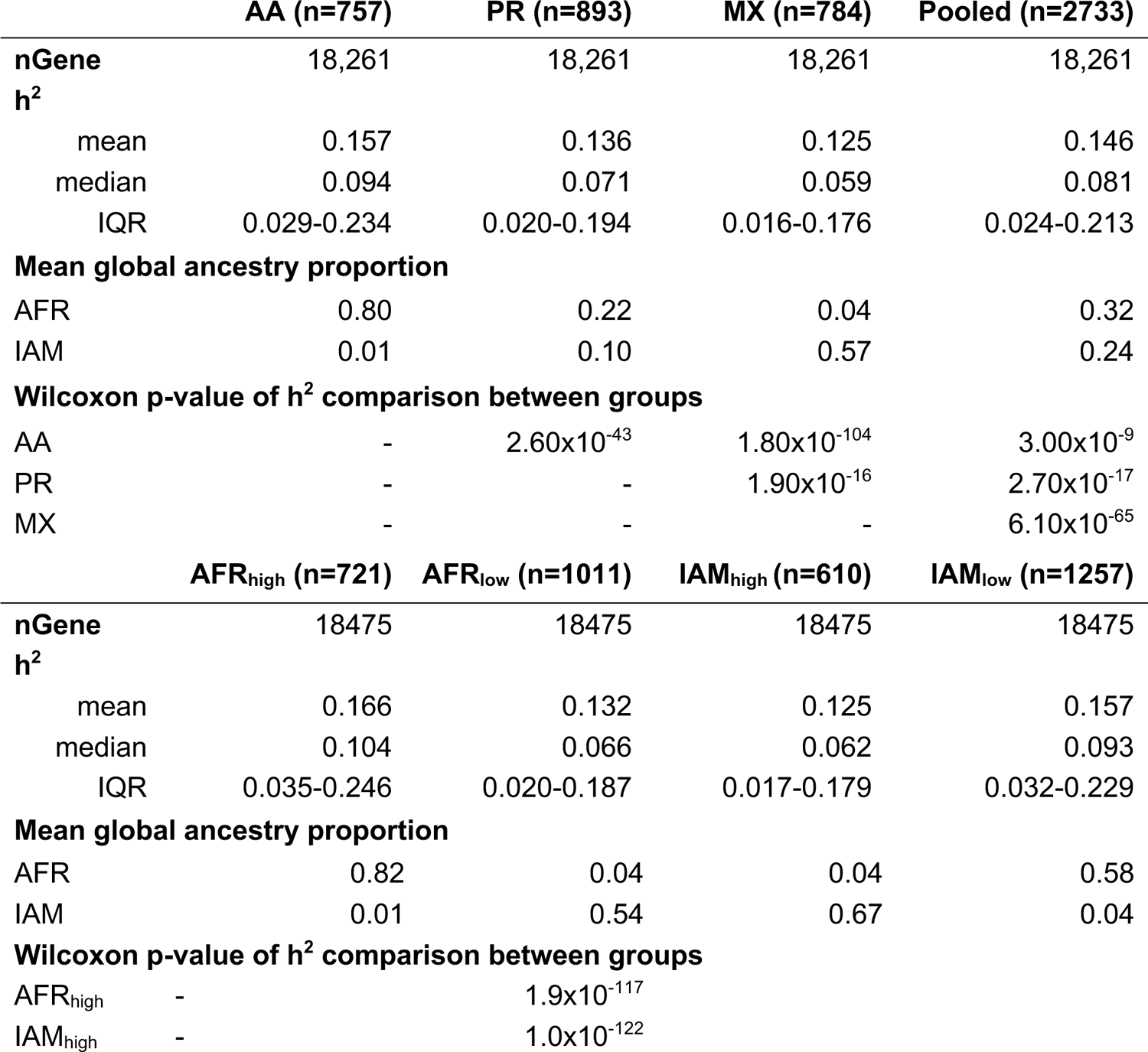
*Cis*-heritability (h^2^) estimated using LDAK-Thin. Analyses were restricted to common variants (MAF >= 0.01) in each population within 1MB flanking regions of the transcription start site.

**Table S6:**
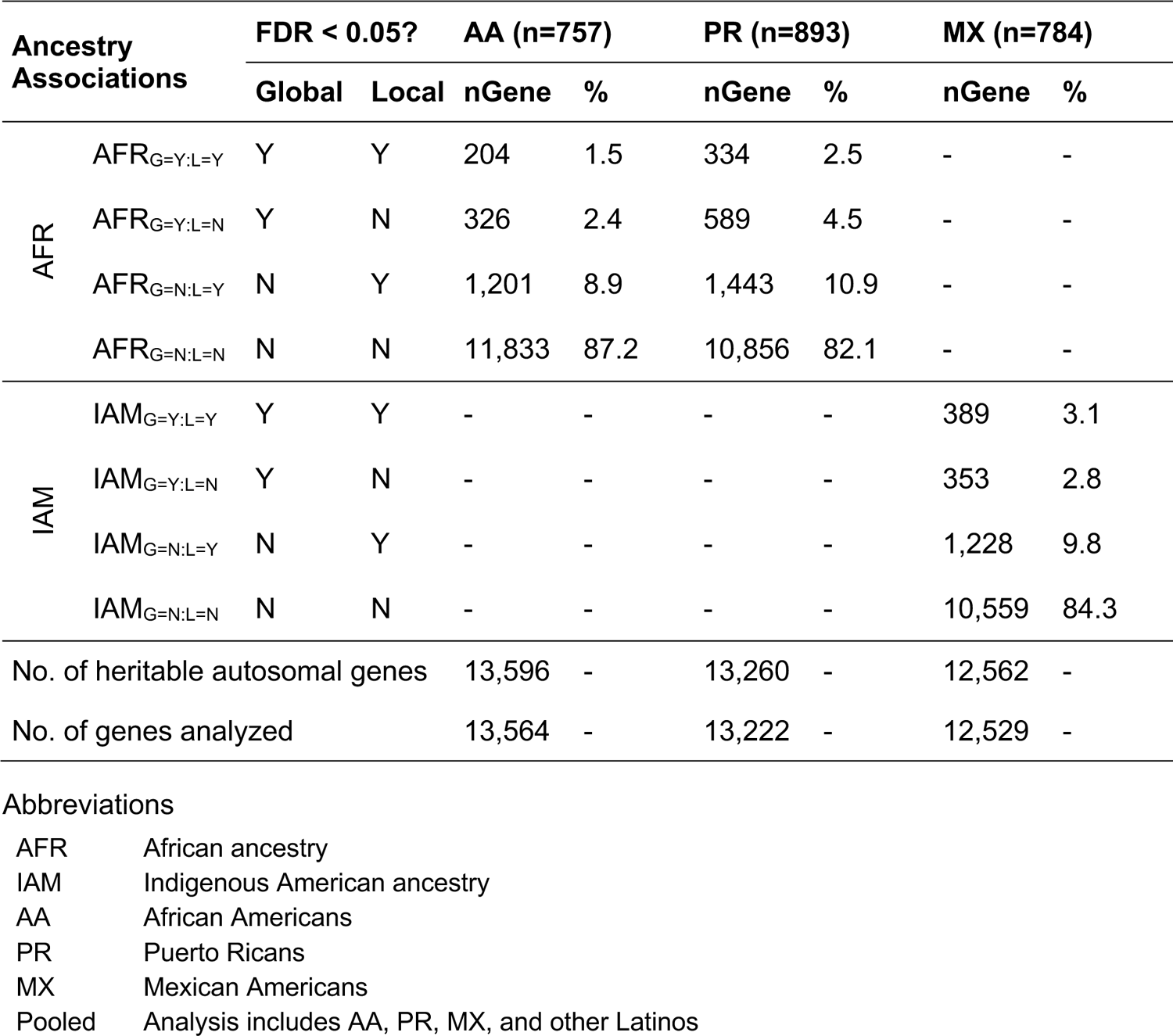
Number of heritable genes significantly associated with global and local ancestry. Analyses were restricted to heritable and autosomal genes with local ancestry estimates, and populations with sufficient variability for a given ancestry comparison. The number of association genes is tabulated for all combinations of global and local ancestry associations. For example, group AFR_G=Y.L=Y_ (global ancestry=Y and local ancestry=Y) includes genes that are associated with both global and local African ancestry at FDR < 0.05 level.

| Ancestry Associations |  | FDR < 0.05? |  | AA (n=757) |  | PR (n=893) |  | MX (n=784) |  |
| --- | --- | --- | --- | --- | --- | --- | --- | --- | --- |
|  |  | Global | Local | nGene | % | nGene | % | nGene | % |
| AFR | $AFR_{G=Y,L=Y}$ | Y | Y | 204 | 1.5 | 334 | 2.5 | - | - |
| | $AFR_{G=Y,L=N}$ | Y | N | 326 | 2.4 | 589 | 4.5 | - | - |
| | $AFR_{G=N,L=Y}$ | N | Y | 1,201 | 8.9 | 1,443 | 10.9 | - | - |
| | $AFR_{G=N,L=N}$ | N | N | 11,833 | 87.2 | 10,856 | 82.1 | - | - |
| IAM | $IAM_{G=Y,L=Y}$ | Y | Y | - | - | - | - | 389 | 3.1 |
| | $IAM_{G=Y,L=N}$ | Y | N | - | - | - | - | 353 | 2.8 |
| | $IAM_{G=N,L=Y}$ | N | Y | - | - | - | - | 1,228 | 9.8 |
| | $IAM_{G=N,L=N}$ | N | N | - | - | - | - | 10,559 | 84.3 |
| No. of heritable autosomal genes |  |  |  | 13,596 | - | 13,260 | - | 12,562 | - |
| No. of genes analyzed |  |  |  | 13,564 | - | 13,222 | - | 12,529 | - |

**Table S7:**
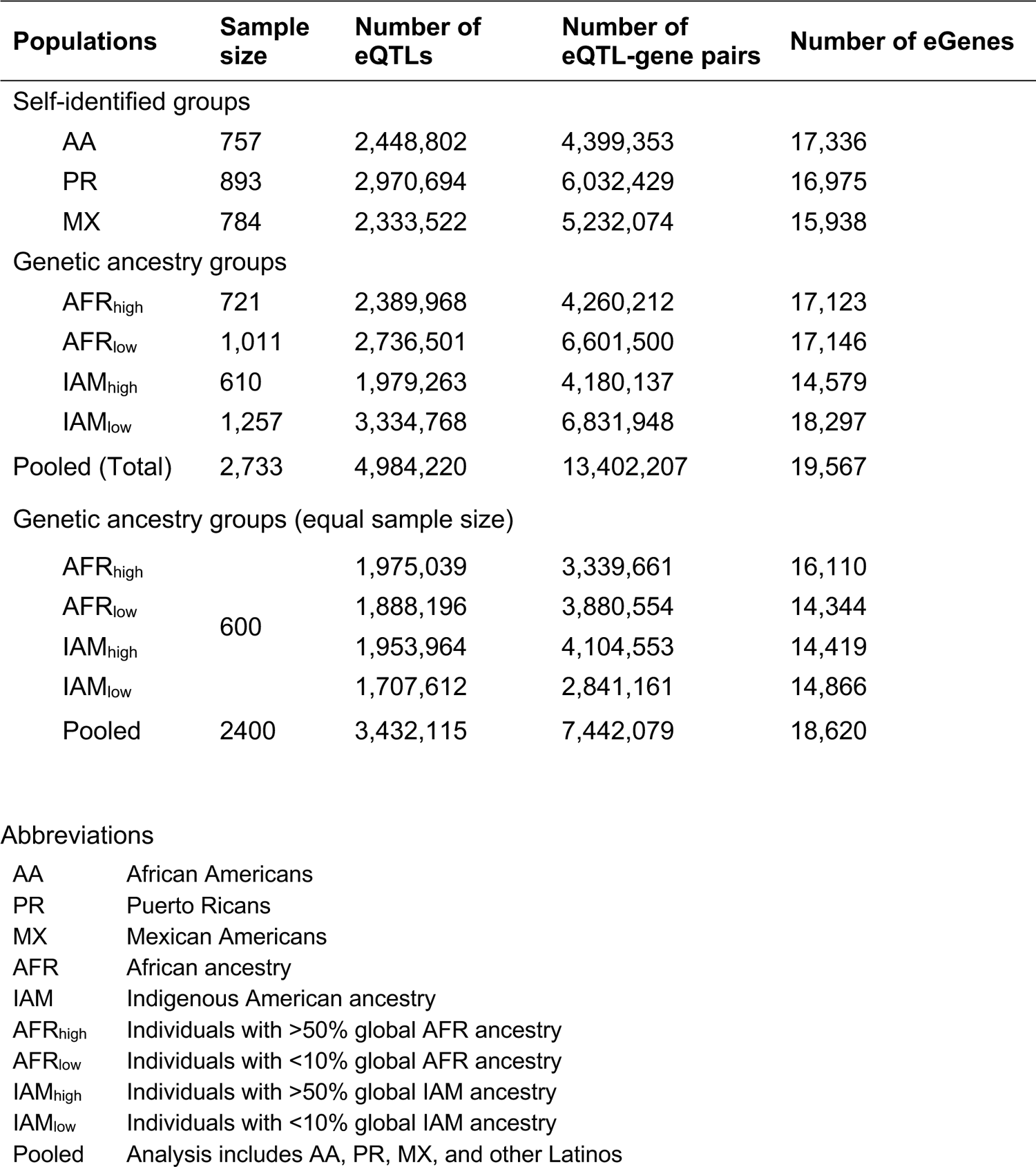
eQTLs and eGenes identified from each population. Results of FastQTL analyses conducted in GALA II / SAGE participants grouped based on self-identified race/ethnicity and genetic ancestry.

| Populations | Sample size | Number of eQTLs | Number of eQTL-gene pairs | Number of eGenes |
| --- | --- | --- | --- | --- |
| Self-identified groups |  |  |  |  |
| AA | 757 | 2,448,802 | 4,399,353 | 17,336 |
| PR | 893 | 2,970,694 | 6,032,429 | 16,975 |
| MX | 784 | 2,333,522 | 5,232,074 | 15,938 |
| Genetic ancestry groups |  |  |  |  |
| AFR<sub>high</sub> | 721 | 2,389,968 | 4,260,212 | 17,123 |
| AFR<sub>low</sub> | 1,011 | 2,736,501 | 6,601,500 | 17,146 |
| IAM<sub>high</sub> | 610 | 1,979,263 | 4,180,137 | 14,579 |
| IAM<sub>low</sub> | 1,257 | 3,334,768 | 6,831,948 | 18,297 |
| Pooled (Total) | 2,733 | 4,984,220 | 13,402,207 | 19,567 |
| Genetic ancestry groups (equal sample size) |  |  |  |  |
| AFR<sub>high</sub> | 600 | 1,975,039 | 3,339,661 | 16,110 |
| AFR<sub>low</sub> |  | 1,888,196 | 3,880,554 | 14,344 |
| IAM<sub>high</sub> |  | 1,953,964 | 4,104,553 | 14,419 |
| IAM<sub>low</sub> |  | 1,707,612 | 2,841,161 | 14,866 |
| Pooled | 2400 | 3,432,115 | 7,442,079 | 18,620 |

|  |  |
| --- | --- |
| AA | African Americans |
| PR | Puerto Ricans |
| MX | Mexican Americans |
| AFR | African ancestry |
| IAM | Indigenous American ancestry |
| AFR<sub>high</sub> | Individuals with >50% global AFR ancestry |
| AFR<sub>low</sub> | Individuals with <10% global AFR ancestry |
| IAM<sub>high</sub> | Individuals with >50% global IAM ancestry |
| IAM<sub>low</sub> | Individuals with <10% global IAM ancestry |
| Pooled | Analysis includes AA, PR, MX, and other Latinos |

**Table S8:**
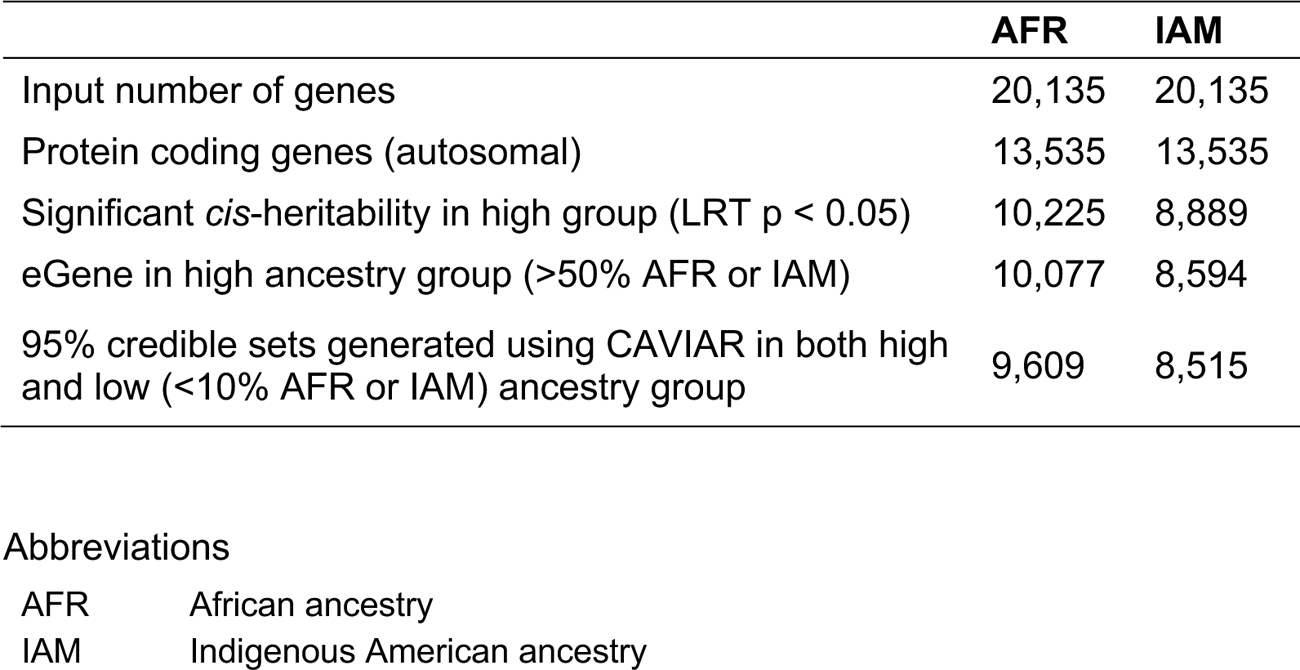
Gene pre-filtering for ancestry-specific eQTL analysis. Significant *cis*-heritability, statistical significance of heritability estimates was determined using LRT p-value provided by GCTA. A total of 9609 and 8515 genes were used as the input to the ancestry-specific eQTL filtering pipeline.

|  | **AFR** | **IAM** |
| --- | --- | --- |
| Input number of genes | 20,135 | 20,135 |
| Protein coding genes (autosomal) | 13,535 | 13,535 |
| Significant *cis*-heritability in high group (LRT p < 0.05) | 10,225 | 8,889 |
| eGene in high ancestry group (>50% AFR or IAM) | 10,077 | 8,594 |
| 95% credible sets generated using CAVIAR in both high and low (<10% AFR or IAM) ancestry group | 9,609 | 8,515 |

|  |  |
| --- | --- |
| AFR | African ancestry |
| IAM | Indigenous American ancestry |

**Table S9:**
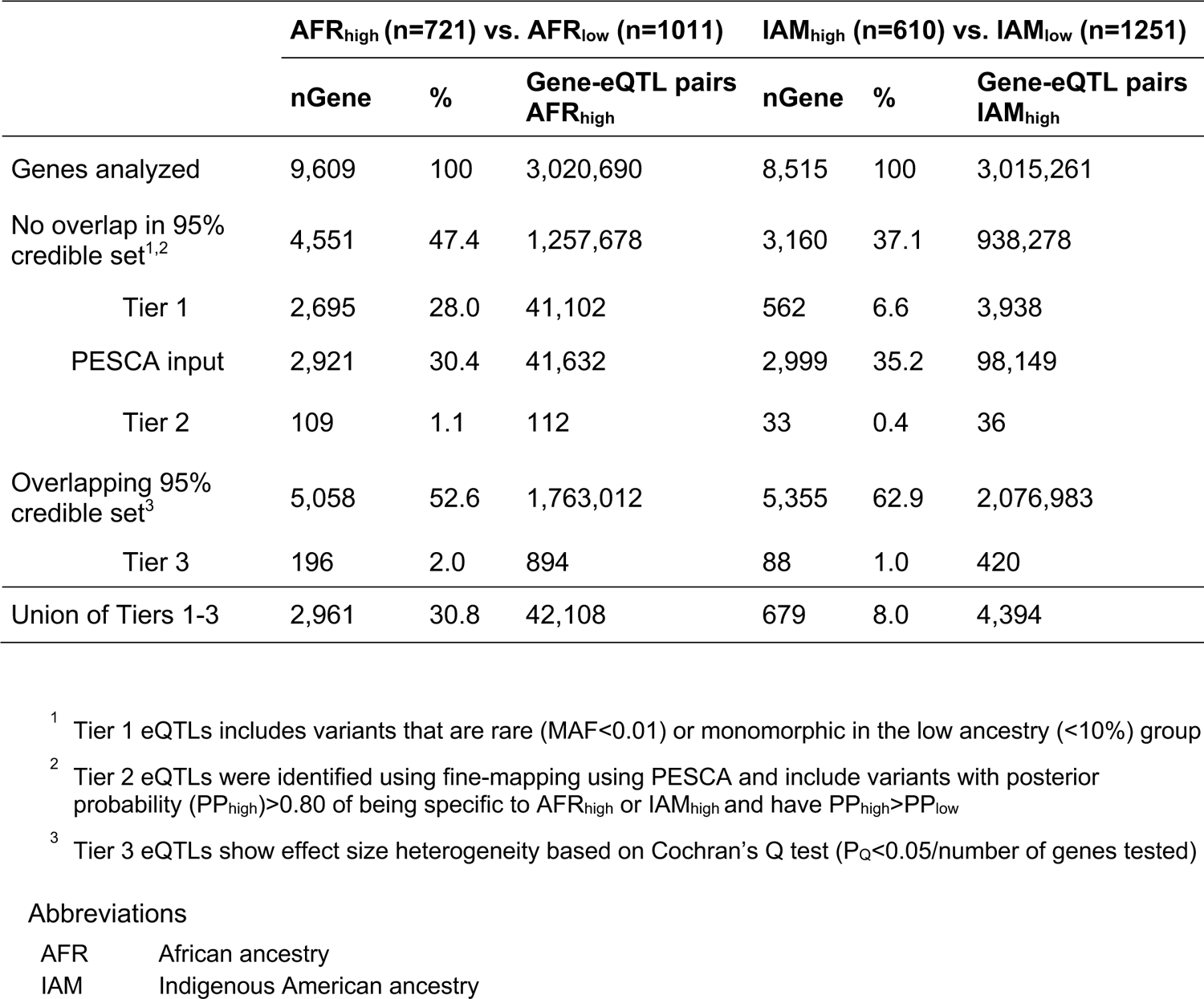
Classification of ancestry-specific eQTLs (anc-eQTLs) using 50% global ancestry cutoff. Analyses were restricted to heritable genes described in Table S8. Comparisons were conducted using >50% as the cut-off for AFR_high_ and IAM_high_ groups. Tier 1 represents the most ancestry-specific eQTL class, followed by Tier 2 anc-eQTLs. Tier 3 eQTLs were detected within overlapping 95% credible sets that are shared between ancestry groups and represent the least ancestry-specific class.

**Table S10:** Classification of ancestry-specific eQTLs (anc-eQTLs) using 70% global ancestry as cutoff. Analyses were restricted to heritable genes described in Table S8. Comparisons were conducted using >70% as the cut-off for AFR_high_ and IAM_high_ groups. Tier 1 represents the most ancestry-specific eQTL class, followed by Tier 2 anc-eQTLs. Tier 3 eQTLs were detected within overlapping 95% credible sets that are shared between ancestry groups and represent the least ancestry-specific class.

|  | AFR<sub>High</sub> (n=653) vs. AFR<sub>Low</sub> (n=1011) |  |  | IAM<sub>High</sub> (n=212) vs. IAM<sub>Low</sub> (n=1251) |  |  |
| --- | --- | --- | --- | --- | --- | --- |
|  | nGene | % | Gene-eQTL pairs AFR<sub>high</sub> | nGene | % | Gene-eQTL pairs IAM<sub>high</sub> |
| Genes analyzed | 9,267 | 100 | 2,653,736 | 4,587 | 100 | 783,676 |
| No overlap in 95% credible set<sup>1,2</sup> | 4,405 | 45.8 | 1,116,628 | 1,726 | 20.3 | 204,927 |
| Tier 1 | 2,620 | 27.3 | 39,300 | 280 | 3.3 | 2,263 |
| Tier 2 | 111 | 1.2 | 111 | 5 | 0.1 | 5 |
| Overlapping 95% credible set<sup>3</sup> | 4,862 | 50.6 | 1,537,018 | 2,861 | 33.6 | 578,749 |
| Tier 3 | 1 | <0.001 | 1 | 0 | 0 | 0 |
| Union of Tiers 1-3 | 2,701 | 28.1 | 39,412 | 284 | 3.3 | 2,268 |

<sup>1</sup> Tier 1 eQTLs includes variants that are rare (MAF<0.01) or monomorphic in the low ancestry (<10%) group
<sup>2</sup> Tier 2 eQTLs were identified using fine-mapping using PESCA and include variants with posterior probability (PP<sub>high</sub>)>0.80 of being specific to AFR<sub>high</sub> or IAM<sub>high</sub> and have PP<sub>high</sub>>PP<sub>low</sub>
<sup>3</sup> Tier 3 eQTLs show effect size heterogeneity based on Cochran's Q test (P<sub>Q</sub><0.05/number of genes tested)

**Table S13:** Trans-eQTL discovery in GALA II/SAGE studies. Independent trans-eQTLs were identified using LD clumping (within 1000 kb windows and LD r^2^<0.1) was performed on trans-eQTLs for each gene. AFR_high_/IAM_high_ groups, individuals with global AFR/IAM ancestry >50%.

| Populations | Sample Size | Trans-eQTLs | Independent trans-eQTLs | eGenes |
| --- | --- | --- | --- | --- |
| AA | 757 | 329 | 39 | 33 |
| PR | 893 | 956 | 67 | 52 |
| MX | 784 | 1,168 | 62 | 51 |
| Pooled | 2,733 | 9,864 | 647 | 414 |
| AFR<sub>high</sub> | 721 | 283 | 36 | 31 |
| Filtered<sup>1</sup> |  | 149 | 26 | 24 |
| IAM<sub>high</sub> | 610 | 691 | 26 | 22 |
| Filtered<sup>2</sup> |  | 350 | 23 | 20 |

<sup>1</sup> All trans-eQTLs detected in AFR<sub>low</sub> group were removed
<sup>2</sup> All trans-eQTLs detected in IAM<sub>low</sub> group were removed

**Table S14:**
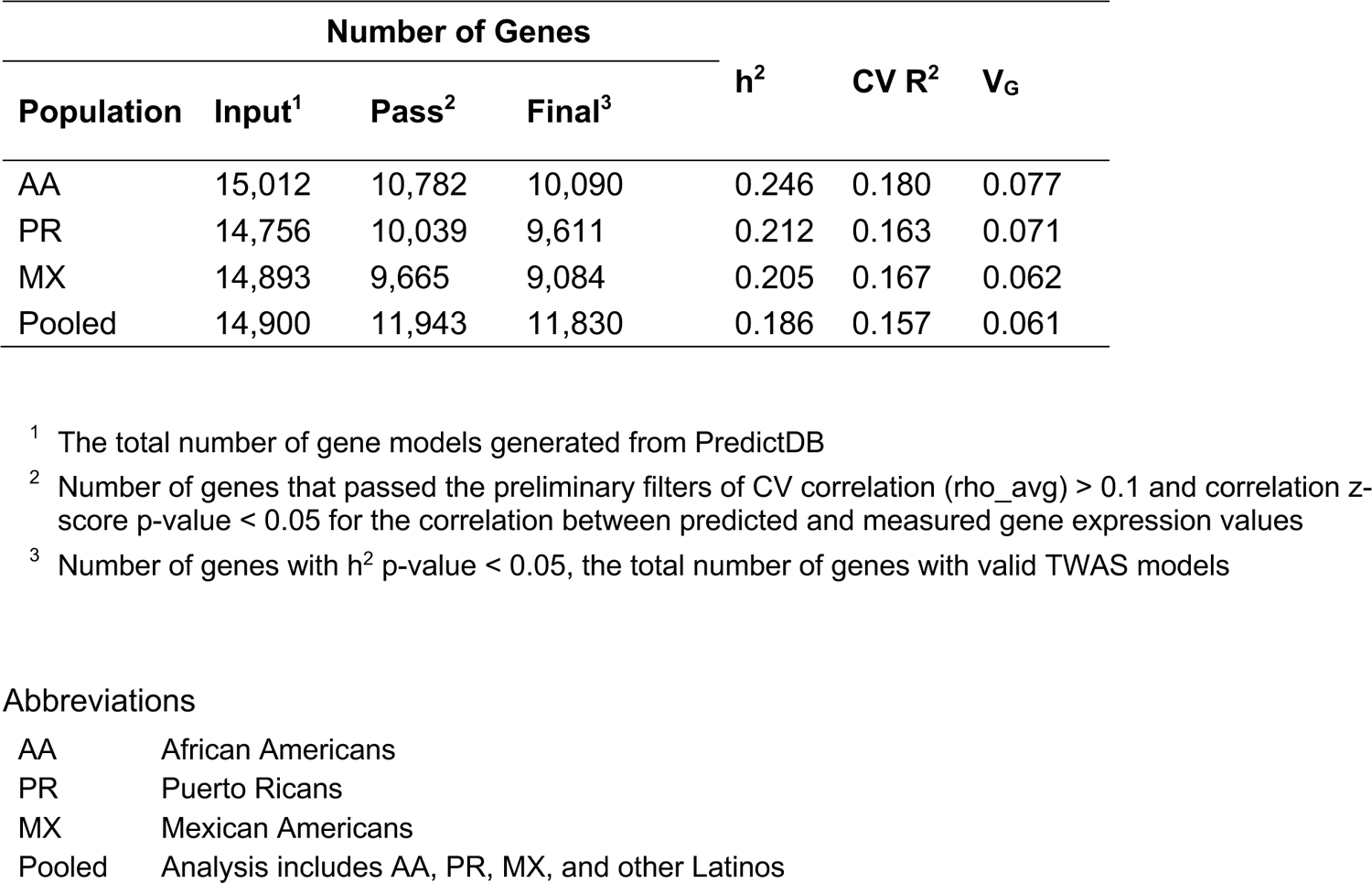
TWAS model performance. Cross-validation (CV) R^2^ of gene expression prediction models generated by PredictDB. Heritability, CV R^2^, and V_G_ are summarized across the final set of genes included in the TWAS models.

| Population | Number of Genes | | | $h^2$ | CV $R^2$ | $V_G$ |
| --- | --- | --- | --- | --- | --- | --- |
|  | Input<sup>1</sup> | Pass<sup>2</sup> | Final<sup>3</sup> |  |  |  |
| AA | 15,012 | 10,782 | 10,090 | 0.246 | 0.180 | 0.077 |
| PR | 14,756 | 10,039 | 9,611 | 0.212 | 0.163 | 0.071 |
| MX | 14,893 | 9,665 | 9,084 | 0.205 | 0.167 | 0.062 |
| Pooled | 14,900 | 11,943 | 11,830 | 0.186 | 0.157 | 0.061 |

<sup>1</sup> The total number of gene models generated from PredictDB
<sup>2</sup> Number of genes that passed the preliminary filters of CV correlation ( $\rho_{\text{avg}} > 0.1$ ) and correlation z-score p-value $< 0.05$ for the correlation between predicted and measured gene expression values
<sup>3</sup> Number of genes with $h^2$ p-value $< 0.05$ , the total number of genes with valid TWAS models

|  |  |
| --- | --- |
| AA | African Americans |
| PR | Puerto Ricans |
| MX | Mexican Americans |
| Pooled | Analysis includes AA, PR, MX, and other Latinos |

**Table S14:** Comparison of TWAS model performance with local ancestry (LA) adjustment. Cross-validation R^2^ of gene expression prediction models with and without local ancestry adjustment. Comparisons were restricted to heritable genes with valid TWAS models.

| Population | Valid TWAS Models | | | CV $R^2$ | |
| --- | --- | --- | --- | --- | --- |
|  | Original | LA-adjusted | Intersection | Original | LA-adjusted |
| AA | 10,090 | 9,848 | 9,701 | 0.186 | 0.177 |
| PR | 9,611 | 9,090 | 8,959 | 0.173 | 0.156 |
| MX | 9,084 | 8,582 | 8,475 | 0.177 | 0.161 |
| Pooled | 11,830 | 11,588 | 11,497 | 0.161 | 0.154 |

**Figure S1:**
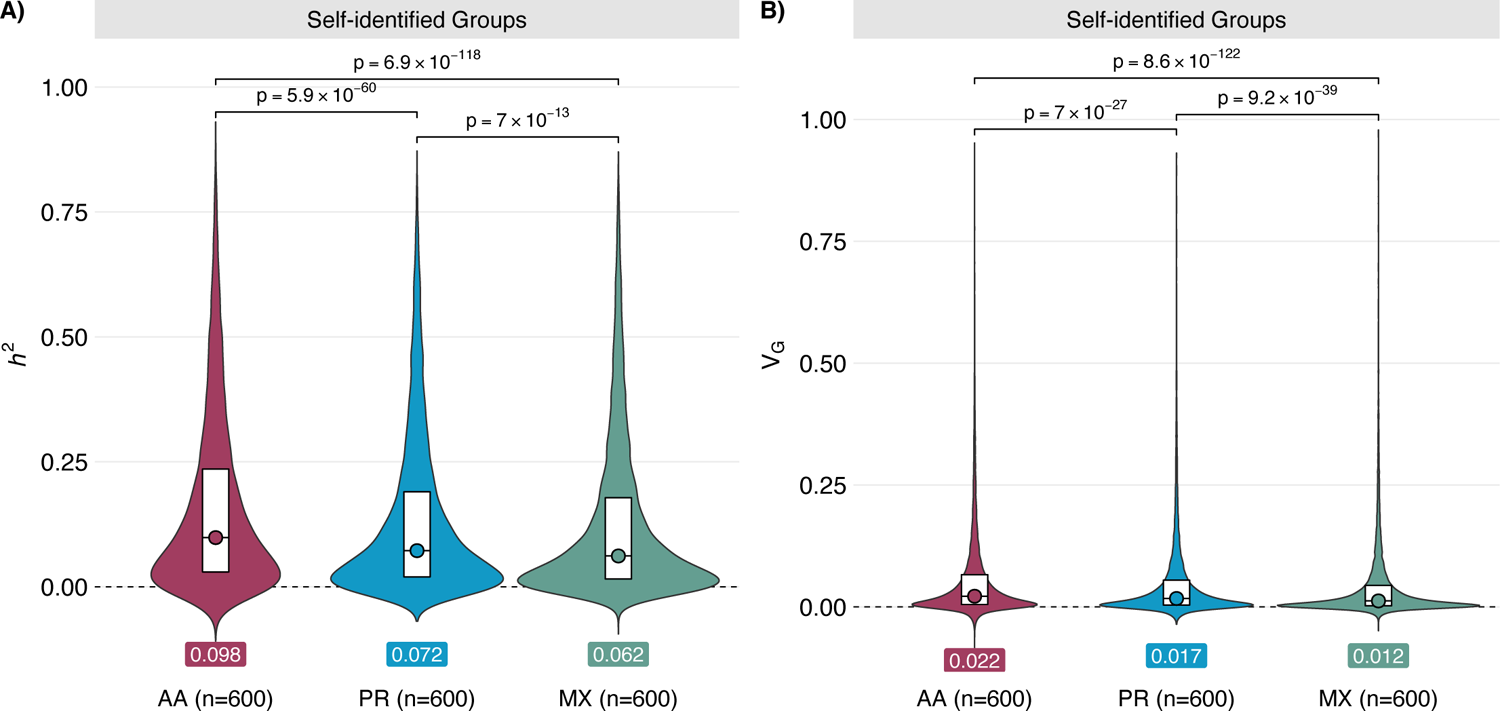
Comparison of *cis*-heritability (*h^2^*) and genetic component of transcriptome variance (V_G_) in a fixed sample size. Within each self-identified race/ethnicity group, individuals were down-sampled to n=600 for all analyses. Median values of h^2^ or V_G_ and two-sided Wilcoxon p-values are annotated.

**Figure S2:**
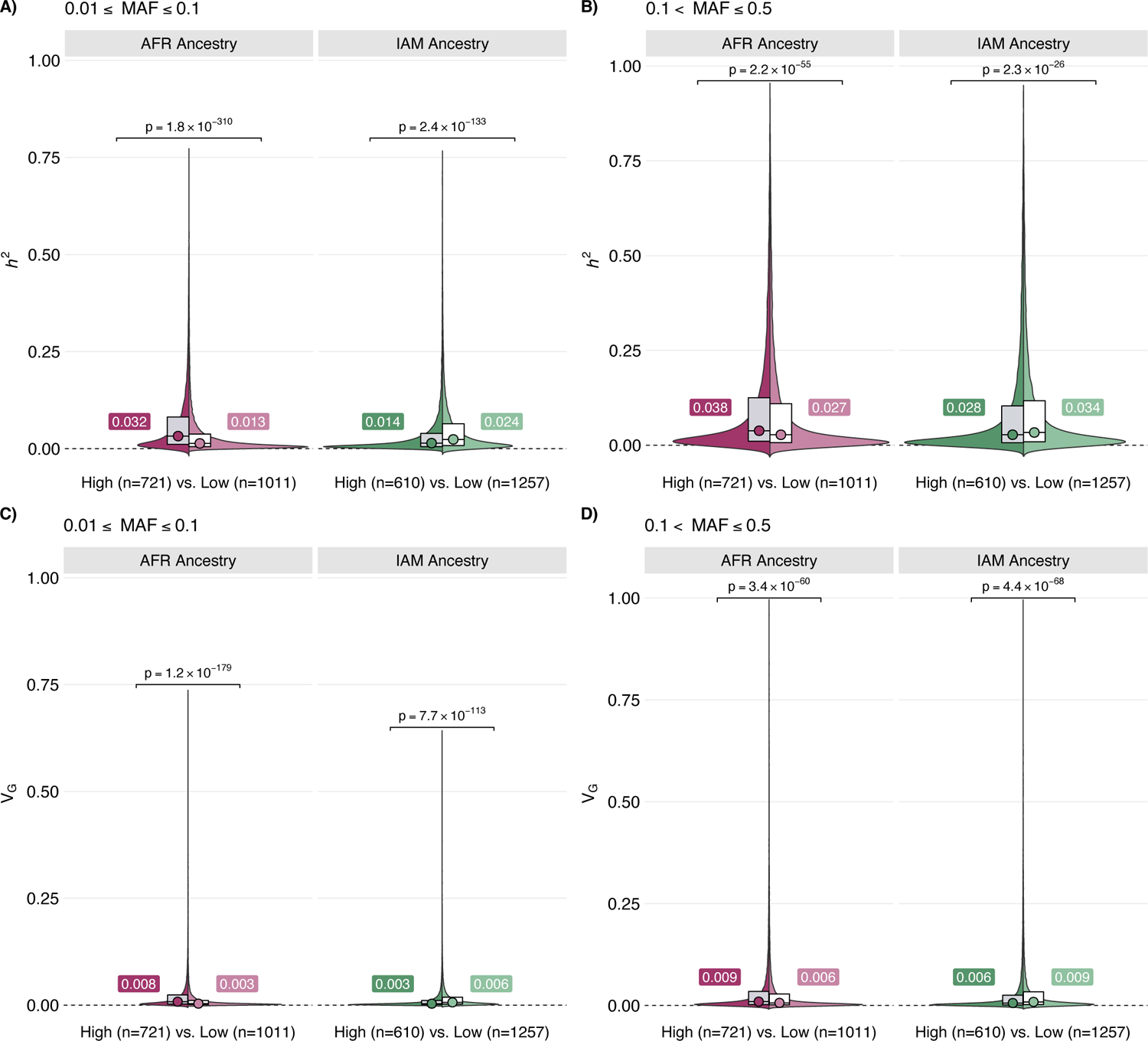
Comparison of *cis*-heritability (*h^2^*) and genetic component of transcriptome variance (V_G_) by genetic ancestry groups and minor allele frequency (MAF). Analyses stratified by genetic ancestry compared individuals with ≥50% global ancestry (High) to participants with <10% of the same ancestry (Low) within each MAF bin. Median values of h^2^ or V_G_ and two-sided Wilcoxon p-values are annotated.

**Figure S3:**
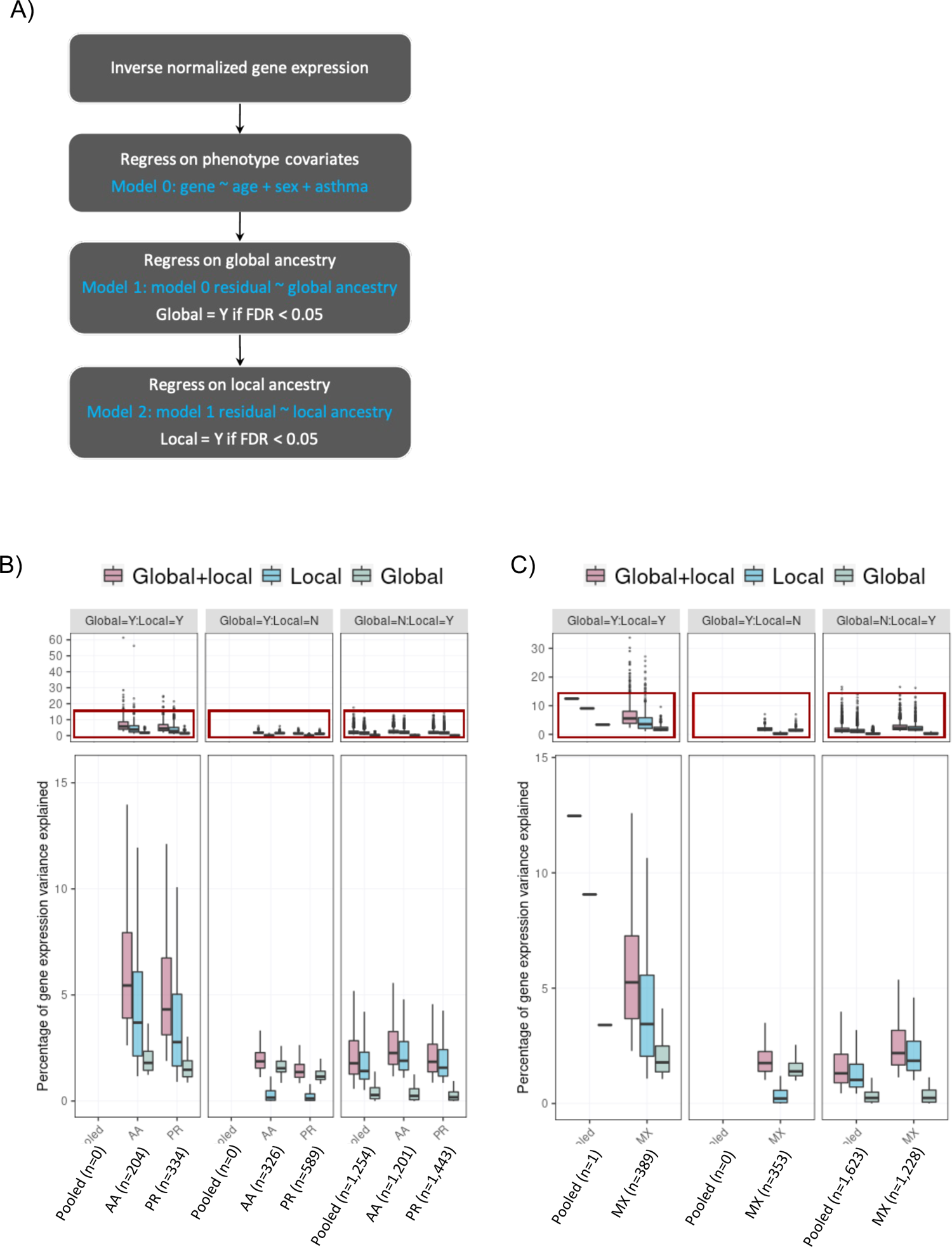
Association of global and local ancestry with gene expression levels. Stepwise local regression was used to identify genes for which global and/or local ancestry had a significant (FDR<0.05) effect on transcript levels. For genes with significant global and/or local ancestry associations, the variance in transcript levels accounted for by African and Indigenous American ancestry. In each panel, inset plots visualize the 0-15% range on the y-axis, without outliers while the full range percentage variance explained are shown in the top panel. Red box highlights the zoomed region as shown in the bottom panel.

**Figure S4:**
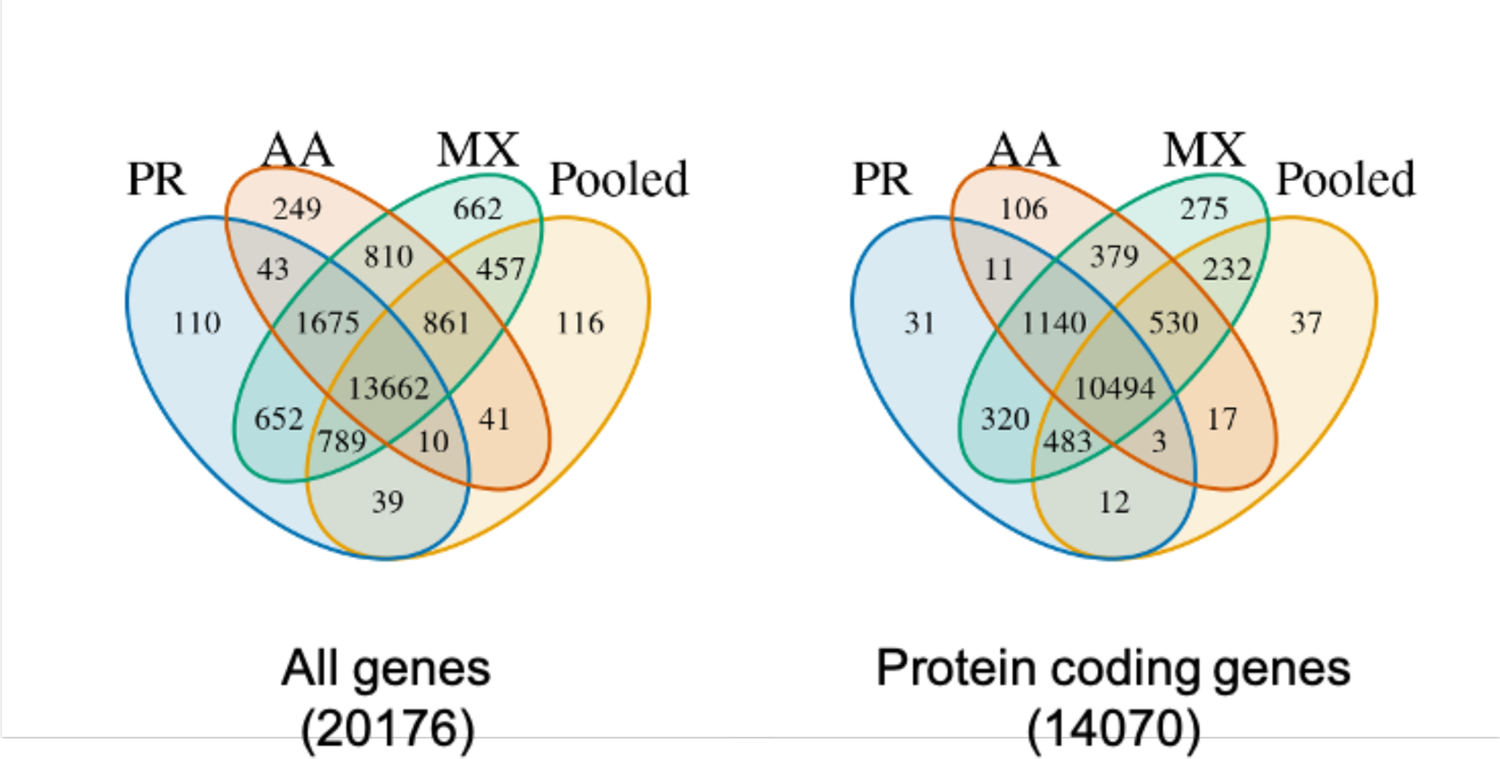
Overlap of eGenes between self-identified race/ethnicity groups

**Figure S5:**
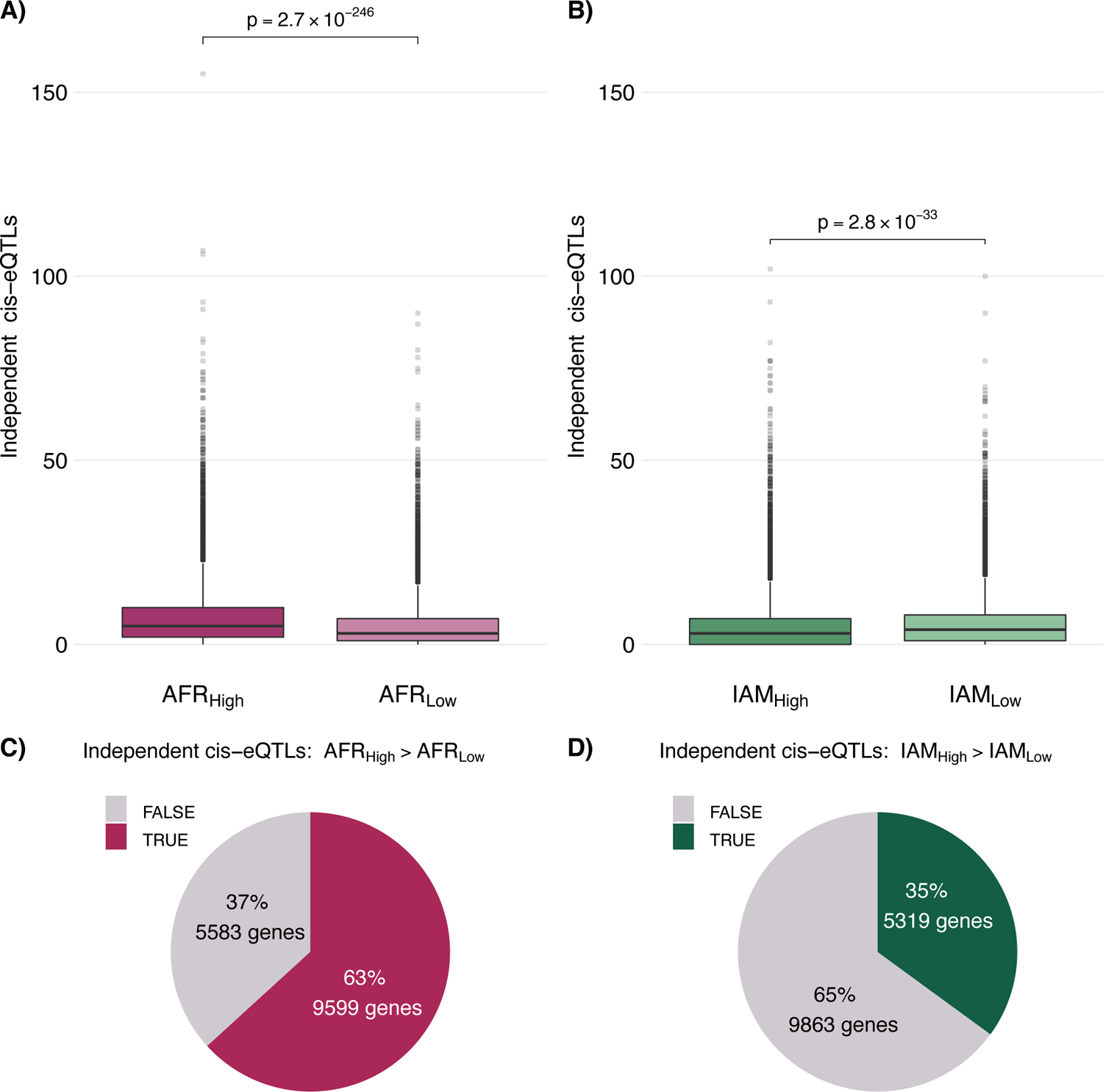
Comparison of independent cis-eQTLs. Sample size was fixed to n=600 for eQTL mapping analyses in each ancestry group. Independent cis-eQTLs were identified by performing LD-based clumping (*r*^2^<0.10) of statistically significant results within each ancestry group. Differences in the distribution of independent cis-eQTLs per gene between AFR_high_ and AFR_low_ **A)** and IAM_high_ and IAM_low_ **B)** ancestry groups were tested using a two-sided Wilcoxon test. Pie charts visualize the proportion of genes with a greater number of cis-eQTLs in AFR_high_ compared to AFR_low_ **C)** and IAM_high_ compared to IAM_low_ **D)**.

**Figure S6:**
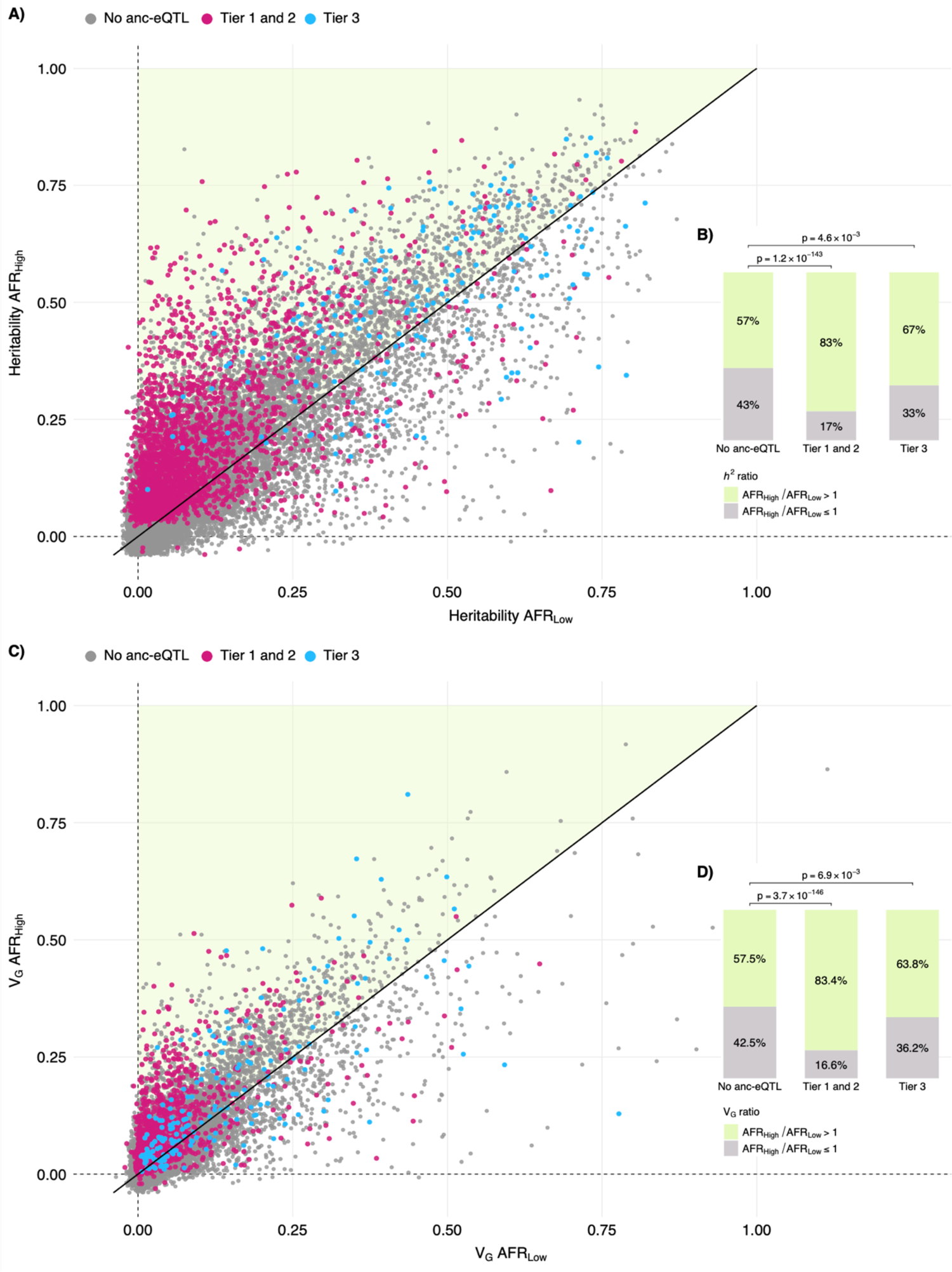
Scatter plots comparing *h^2^* and V_G_ by African ancestry. Estimates of h^2^ and V_G_ for each gene are compared for individuals with ≥50% global African ancestry (AFR_High_) to participants with <10% AFR ancestry (AFR_Low_). Genes containing ancestry-specific eQTLs are are highlighted. The proportion of genes falling off the diagonal, with higher h_2_ or V_G_ in AFR_High_ than AFR_Low_, is visualized and compared using a two-sided binomial test.

**Figure S7:**
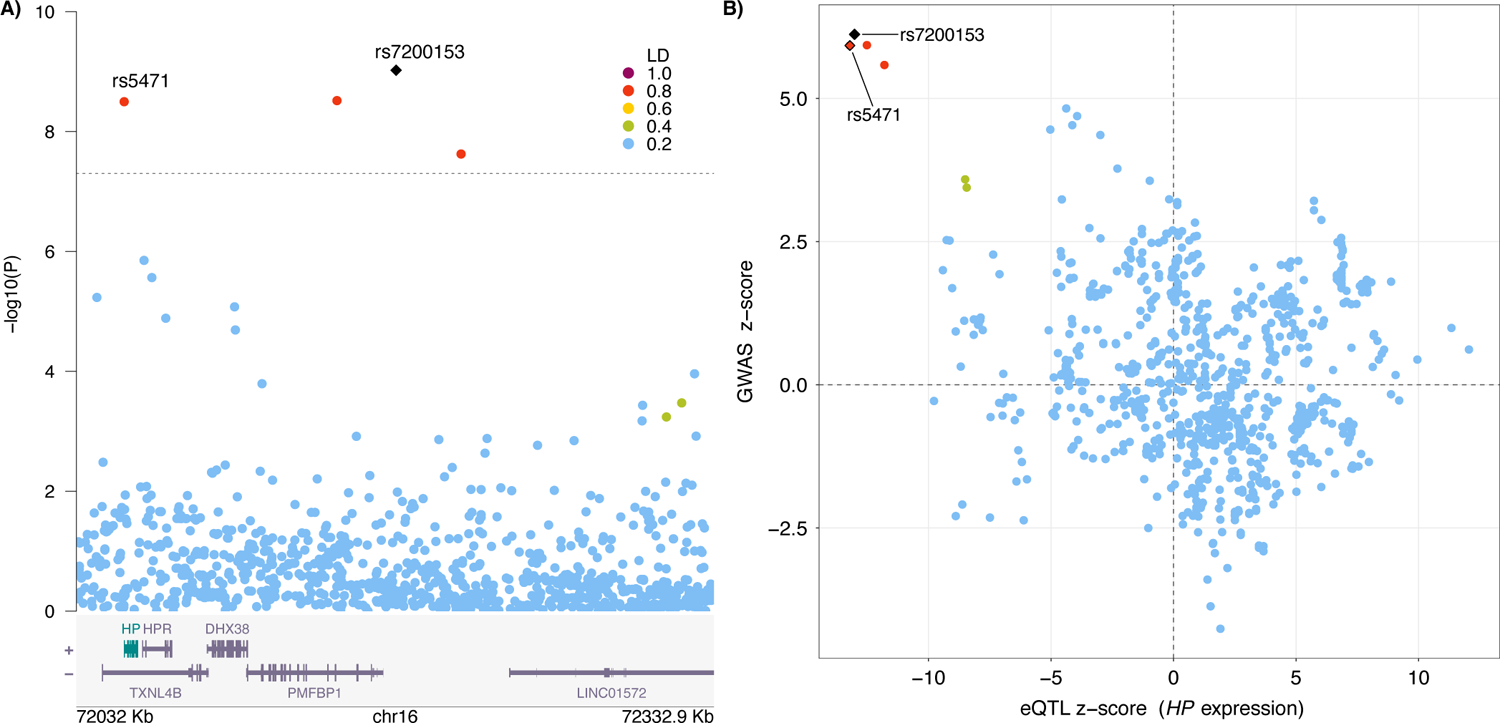
Colocalization of haptoglobin (*HP*) expression and total cholesterol. We observed strong evidence of colocalization, with posterior probability (PP)=0.997, between GALA/SAGE eQTLs for *HP* and GWAS summary statistics from PAGE for total cholesterol. The 95% credible set contained two variants: rs7200153 (PP_SNP_=0.519) and rs5471 (PP_SNP_=0.481). Each plot shows variants colored based on LD with respect to rs7200153, which had the lowest GWAS p-value in PAGE.

**Figure S8:**
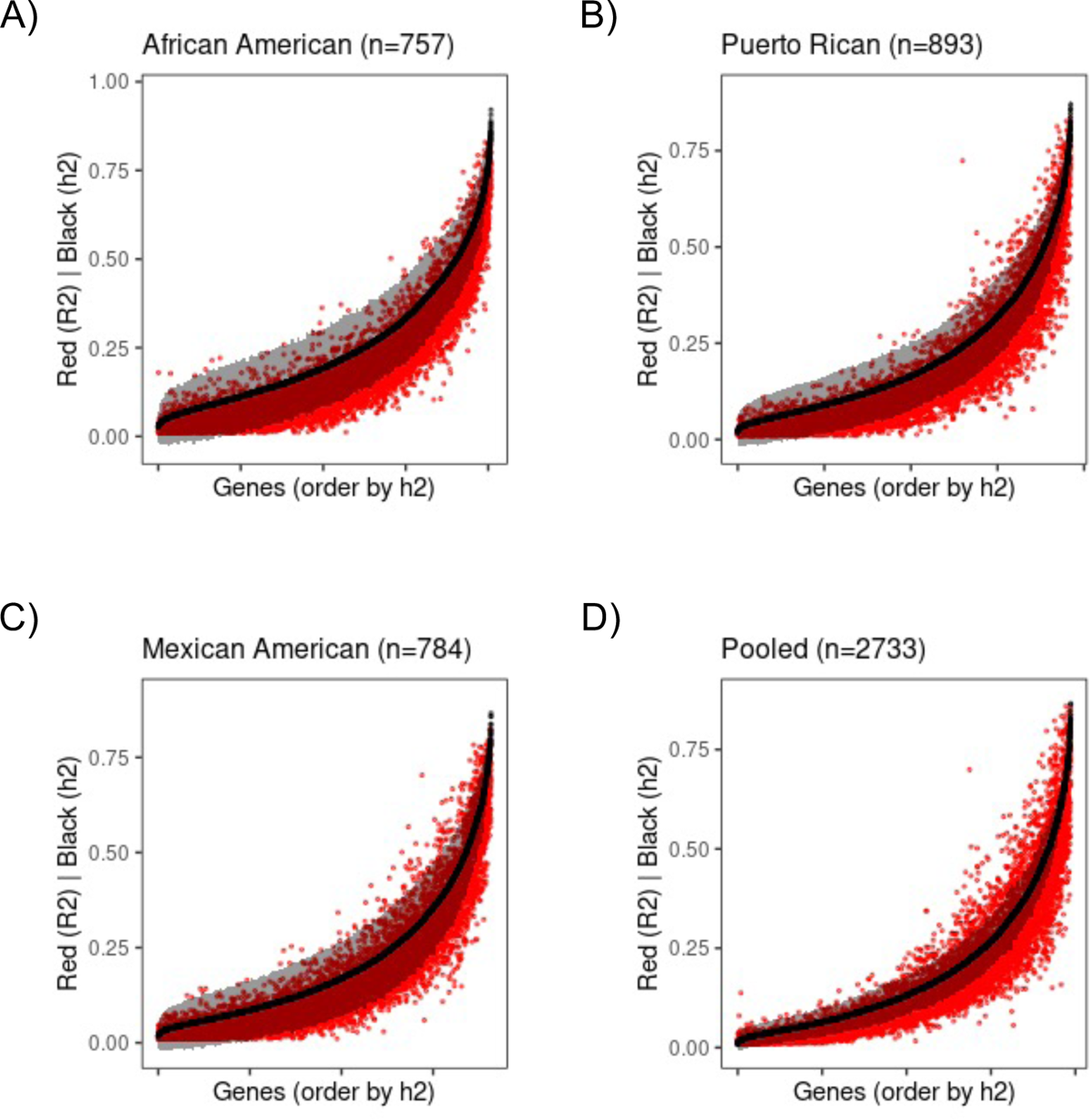
Cross validation R^2^ of gene prediction models generated from PredictDB. Cross validation (CV) R^2^ of each gene expression prediction model is represented by a red dot. Cis-heritability of the gene, represented as black dot with 95% confidence interval in grey, was shown to indicate upper bound of CV R^2^. Genes are sorted in ascending order of h^2^.

**Figure S9:**
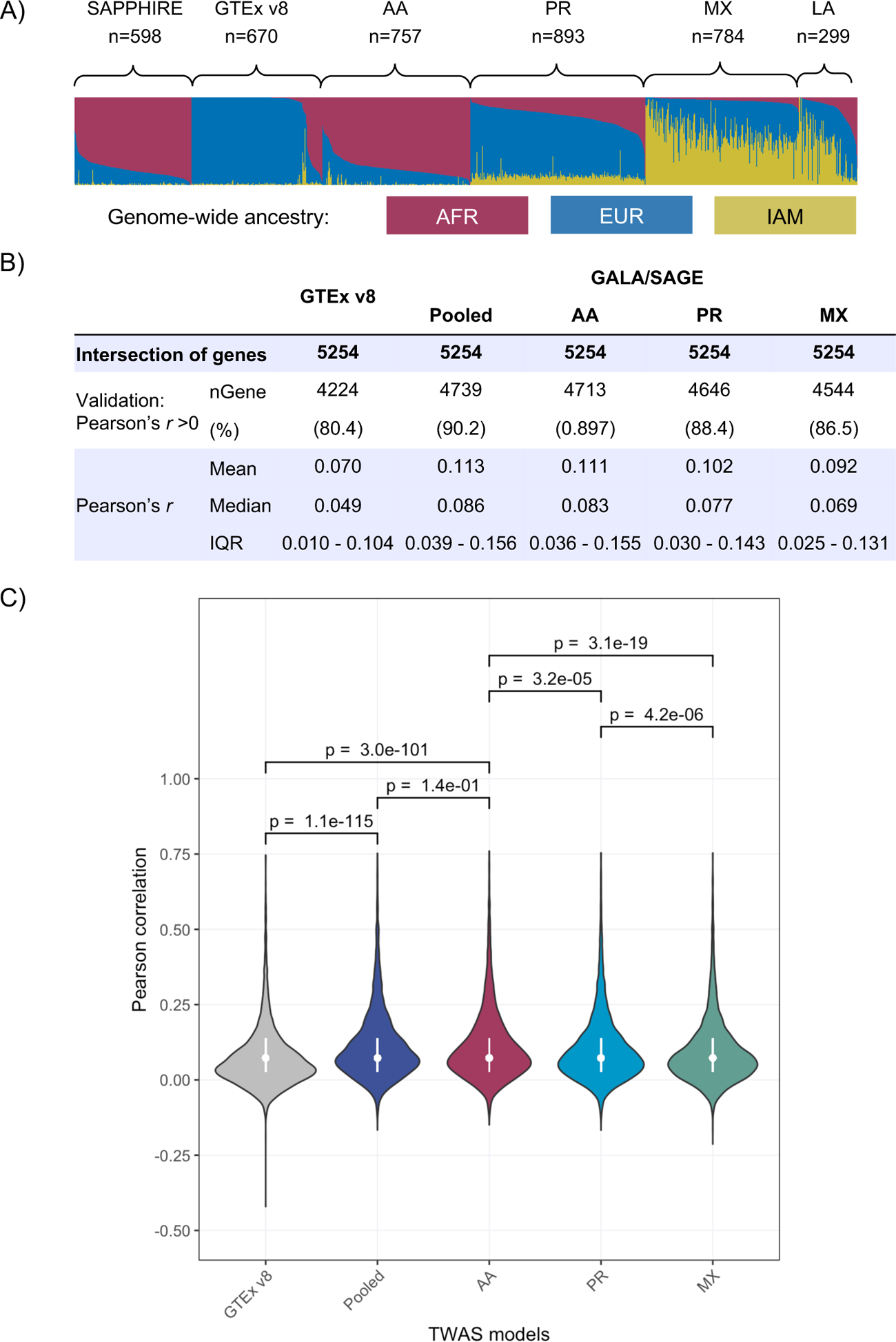
Out of sample validation of TWAS models in the SAPPHIRE. Admixture plots for the SAPPHIRE validation study and each of the training samples used to develop the TWAS models are shown in panel A). Validation results are shown for the subset of genes (n=5254) that were available in GTEx and GALA/SAGE models. Correlation between the predicted and measured gene expression levels is summarized in panel B) and the full distribution of correlation coefficients is shown in C).

**Figure S10:**
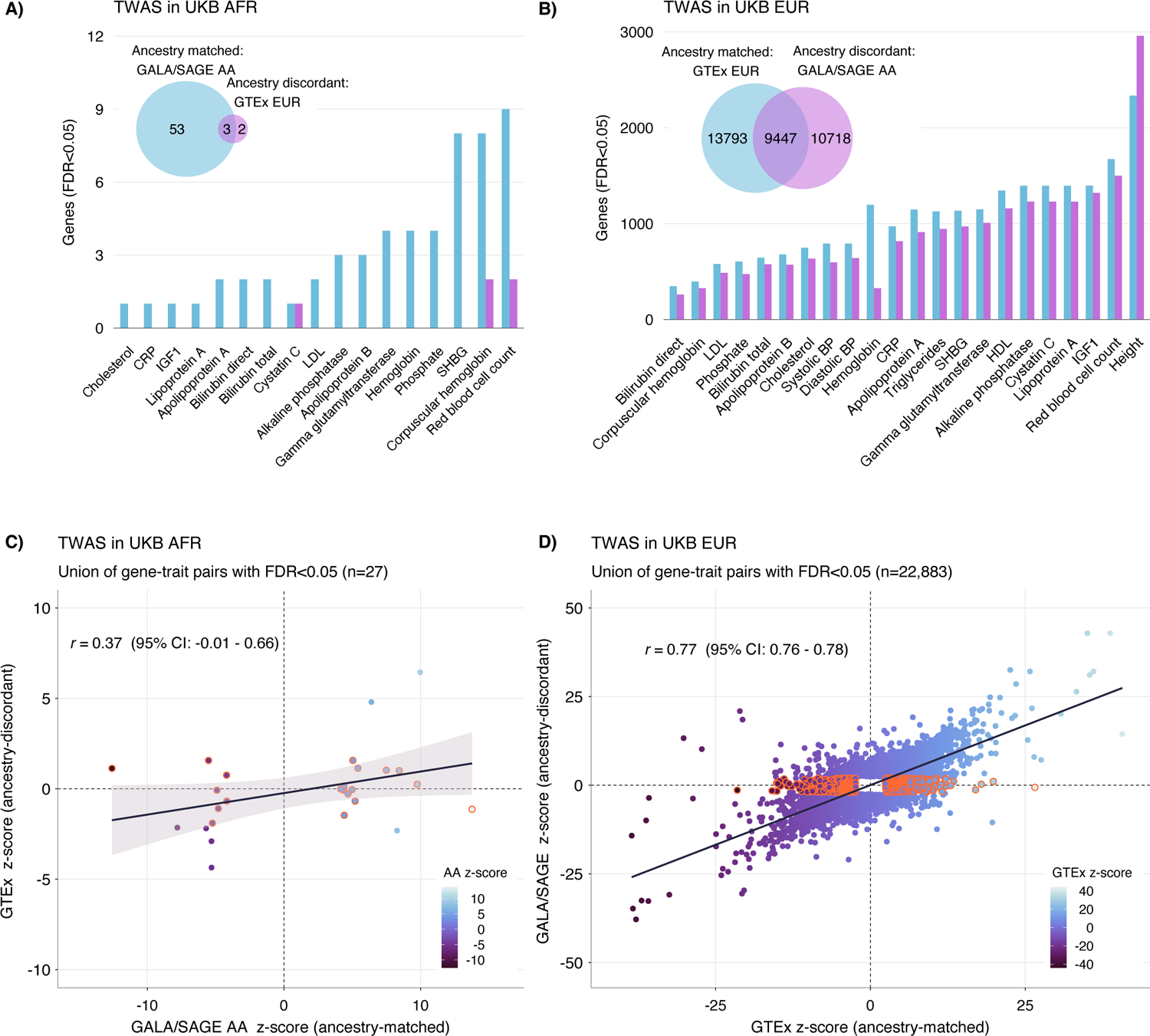
Summary of TWAS results in UK Biobank (UKB). Comparative TWAS analyses in UKB were conducted using GTEx v8 whole blood models and GALA/SAGE models trained in African Americans (AA). Number of associated genes in ancestry matched and ancestry discordant analyses is summarized in for UKB European (EUR) ancestry subjects in **A)** and UKB African (AFR) ancestry subjects in **B)**. Correlation between the z-scores for statistically significant findings in UKB EUR **C)** and UKB AFR **D)** are shown for genes that were present in both models. Genes highlighted in orange had FDR<0.05 using ancestry-matched models but did not reach nominal significance (TWAS p-value>0.05) using ancestry discordant models.

**Figure S11:**
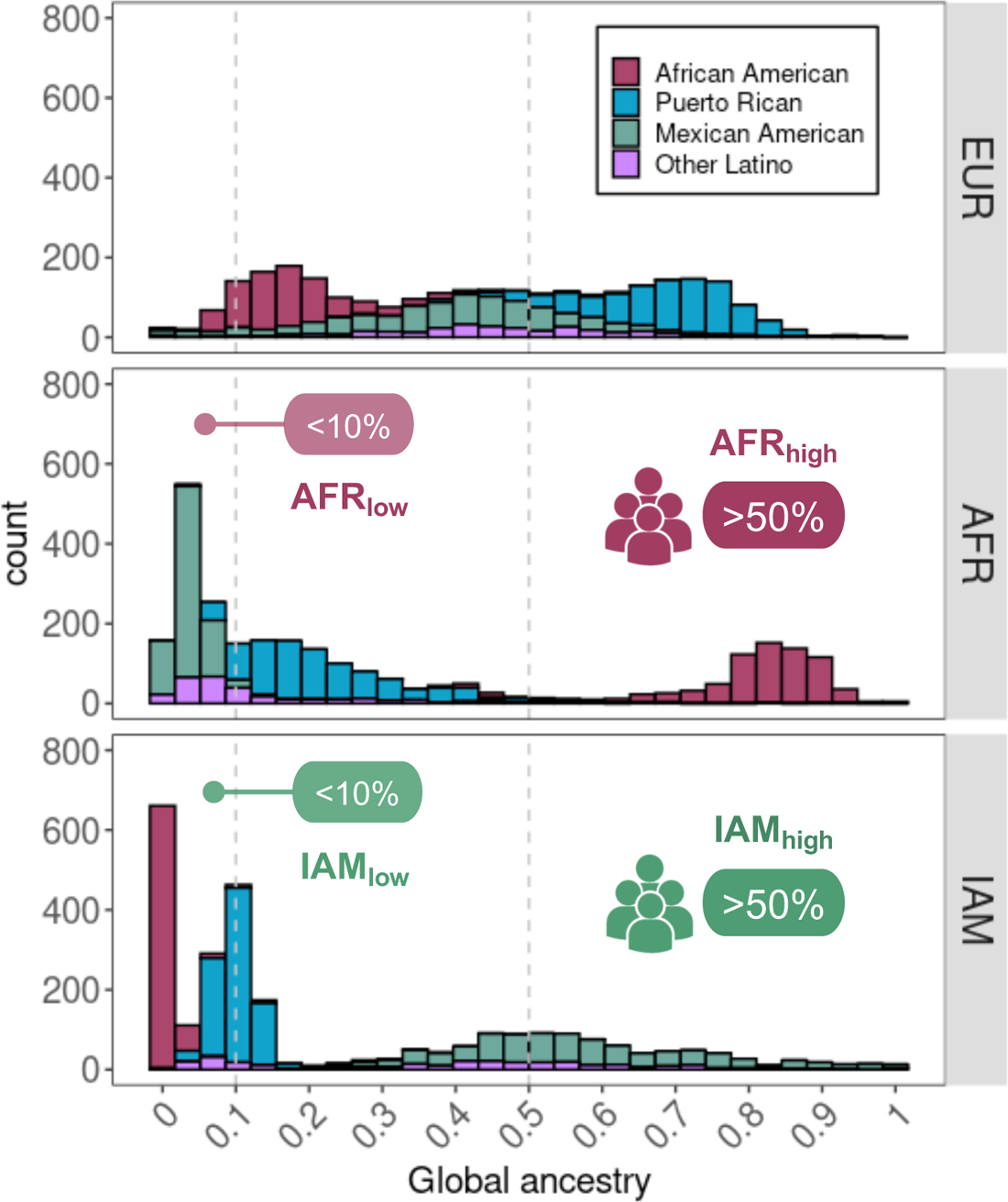
Distribution of global genetic ancestry in GALA/SAGE participants.

**Figure S12:**
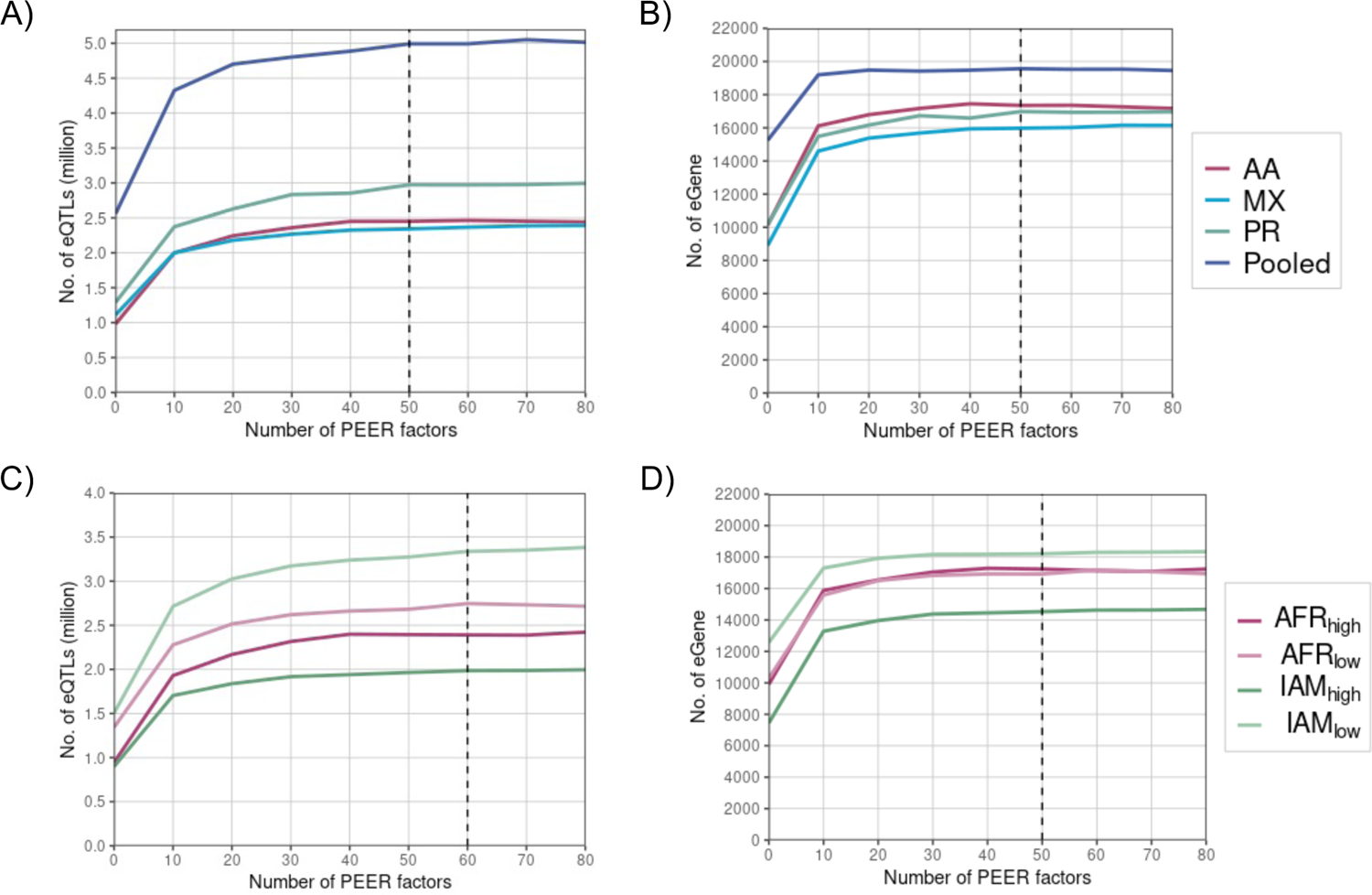
Selection of PEER factors for downstream analysis. Each panel visualizes the number of eQTLs and eGenes identified using different number of PEER factors included as covariates. Vertical dashed lines indicate the number of PEER factors selected for the final analysis with the goal of maximizing eQTL discovery.

